# A novel Bayesian factor analysis method improves detection of genes and biological processes affected by perturbations in single-cell CRISPR screening

**DOI:** 10.1101/2022.02.13.480282

**Authors:** Yifan Zhou, Kaixuan Luo, Mengjie Chen, Xin He

## Abstract

CRISPR screening coupled with single-cell RNA-sequencing has emerged as a powerful tool to characterize the effects of genetic perturbations on the whole transcriptome at a single-cell level. However, due to the sparsity and complex structure of data, analysis of single-cell CRISPR screening data remains challenging. In particular, standard differential expression analysis methods are often under-powered to detect genes affected by CRISPR perturbations. We developed a novel method for such data, called Guided Sparse Factor Analysis (GSFA). GSFA infers latent factors that represent co-regulated genes or gene modules, and by borrowing information from these factors, infers the effects of genetic perturbations on individual genes. We demonstrated through extensive simulation studies that GSFA detects perturbation effects with much higher power than state-of-the-art methods. Using single-cell CRISPR data from human CD8^+^ T cells and neural progenitor cells, we showed that GSFA identified biologically relevant gene modules and specific genes affected by CRISPR perturbations, many of which were missed by existing methods, providing new insights into the functions of genes involved in T cell activation and neurodevelopment.

## Introduction

The discovery of Clustered Regularly Interspaced Short Palindromic Repeats (CRISPR) and subsequent development of the CRISPR-Cas9 system for genomic editing have revolutionized many fields of biology^1,2^. A powerful application of the CRISPR-Cas9 system is pooled CRISPR screening, where many genes or genomic sites are edited at the same time to screen for genes with certain functions or phenotypes. This approach has enabled the identification of many genes involved in processes such as cell proliferation and survival, immune responses, and drug resistance^3–5^. Technologies such as CROP-seq^6^ and Perturb-seq^7^ have further improved the power of CRISPR by combining multiplexed CRISPR screening with single-cell RNA-seq (scRNA-seq). By linking guide RNAs (gRNAs) to single cell transcriptomes, these technologies provide comprehensive molecular readouts of the target perturbations within single cells. Single-cell CRISPR screening technologies have found many applications, for example, in studies of developmental regulators, genes involved in immune responses, and regulatory elements involved in human diseases^8–11^.

Nevertheless, the analysis of single-cell CRISPR screening data remains challenging. A routine strategy for analyzing gene expression data is differential expression analysis, whereby the effects of genetic perturbation on the transcriptome are assessed one gene at a time^12,13^. When applied to single cell screening data, however, this type of analysis can be under-powered due to the sparsity and noise inherent to scRNA-seq data, as well as the relatively small numbers of cells (often hundreds) per perturbation in typical experiments. Another commonly used analysis is to cluster cells based on their transcriptome similarity, and then assess whether cells with a specific perturbation are enriched or depleted in any cluster^10,14^. However, this clustering-based approach has several limitations. CROP-seq studies often use relatively homogenous populations of cells to minimize variation across cells and increase the power of discovering transcriptional effects; thus, it can be challenging to partition cells into distinct clusters. Furthermore, the clustering approach does not explicitly link the perturbations with the affected genes, limiting our understanding of the downstream effects of the perturbations. Given the limitations of standard differential expression and clustering-based analyses, rigorous statistical methods that accommodate the unique features and complexities of single-cell CRISPR screening data are greatly needed.

Our proposed approach is motivated by the observation that genetic perturbations typically affect expression, not one gene at a time, but many related genes simultaneously. Indeed, transcriptomes often vary across cell types, conditions, and individuals in a modular fashion, reflecting the underlying coordinated regulation of genes in the same or related pathways. These gene modules can be inferred by matrix factorization and related techniques^15–22^. We propose to infer gene modules from scRNA-seq data, and borrow information across genes to improve the power of detecting differentially expressed genes. Existing factor analysis methods, however, are not readily applied to single-cell CRISPR screening data, as the factors are not directly linked with genetic perturbation, and the effects of perturbation on the expression of individual genes are not assessed.

Here we present Guided Sparse Factor Analysis (GSFA), a framework for analyzing single-cell CRISPR screening data that bridges factor analysis and differential expression analysis. GSFA assumes the effects of genetic perturbations are mediated through a set of gene modules. These gene modules, representing biological pathways or functional units, are captured mathematically as latent factors. GSFA evaluates associations of the genetic perturbations with these latent factors, providing information on the module-level effects of the perturbations. Compared with single-gene level differential expression analysis, this factor association analysis may be more sensitive. Indeed, expression of a single gene is noisy, and is influenced by potentially many sources. In contrast, latent factors represent the main dimensions of variation of many genes, and can be thought of as “denoised” versions of gene expression, possibly improving the detection of the effects of perturbations. While our approach is formulated in terms of latent factors, we can still summarize the effects of a perturbation on individual genes as the sum of effects mediated by all the factors. We benchmarked our method through extensive simulation studies and real data applications. GSFA identifies biologically relevant modules, and has better power to detect differentially expressed genes (DEGs) than alternative methods, providing novel insights into the molecular mechanisms regulating T cell activation and neuronal differentiation.

### Results

### GSFA method overview

GSFA is a Bayesian statistical model that unifies factor analysis and estimation of the effects of target perturbations. The input of GSFA consists of two matrices: a normalized gene expression matrix across cells, and a “perturbation matrix” that records gRNA perturbations in each cell (Fig. 1). GSFA assumes that the perturbation of a target gene affects certain latent factors, which in turn changes the expression of individual genes. These assumptions lead to a two-layer model. In the first layer, the expression matrix (Y) is decomposed into the product of the factor matrix (Z) and the weights of genes on factors (gene loading, W). In the second layer, GSFA captures the dependency of factors (Z) on perturbations (G) via a multivariate linear regression model (Fig. 1).

**Figure 1:**
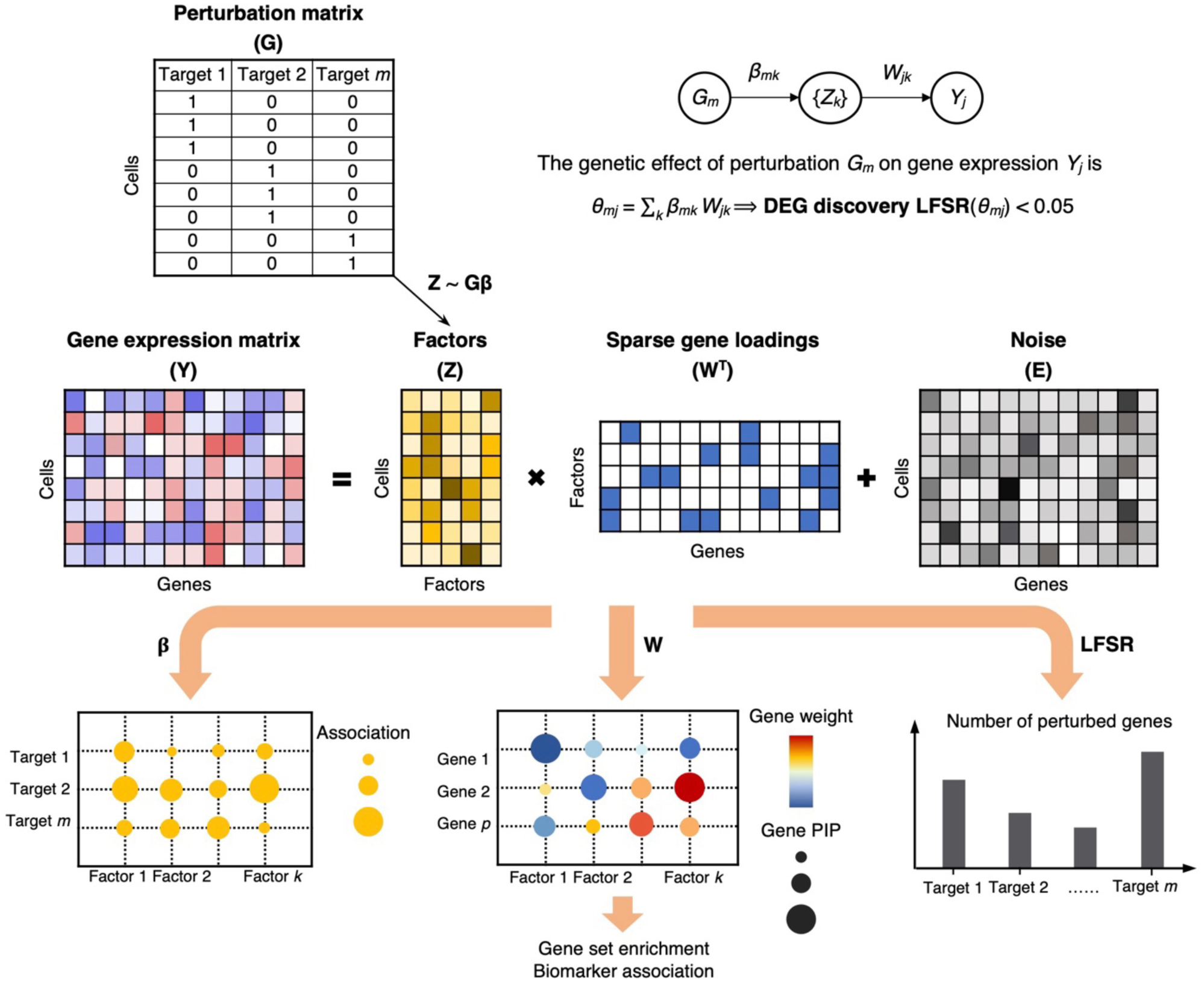
GSFA model and its application on real data. Top: the input of GSFA includes the perturbation matrix and the gene expression matrix; bottom: the output of GSFA includes the effects of perturbations on targets (β), the gene loading matrix (W), and the list of genes affected by each perturbation after LFSR thresholding.

The main unknowns of the model are the values of the factors (Z), the gene loading on factors (W), and the effects of perturbations on the factors (β). We assume a standard normal prior distribution of Z, and a “spike-and-slab” prior of β, assuming that the effects come from either a normal distribution or a point mass at 0^23^. This sparse prior of β encodes the intuition that a genetic perturbation likely affects only a small number of factors. For the gene loading matrix W, we also use a sparse prior to limit the number of genes contributing to a factor, facilitating the biological interpretation of factors. We evaluated two choices, the standard spike-and-slab prior, and a normal-mixture prior, where the effect is sampled from a mixture of two normal distributions, one “foreground” component capturing true effects, and the other a “background” component absorbing small effects^24,25^. This comparison is motivated by a well-known problem in count-based RNA-seq data analysis: because the total read count in a sample is fixed, activation of some genes indirectly reduces the read counts in all other genes, resulting in weakly correlated expression across many genes. Thus, even when a factor affects only a small set of genes, it may appear to be correlated with many other genes, making it hard to infer sparse factors. A normal-mixture prior may help address this problem, because of its “background” component with small effects. Indeed, it shows better results in our simulations compared with the spike-and-slab prior (see below), so is used as our default prior.

We used a Gibbs sampling algorithm to obtain posterior samples of model parameters. For any parameter with a sparse prior, the probability that its value is non-zero or comes from the foreground component, is denoted as posterior inclusion probability (PIP). PIPs quantify whether a perturbation affects a certain factor, or whether a gene has loading on a factor. The factors can then be interpreted, *e*.*g*. through gene ontology (GO) enrichment analysis of genes loaded on the factors, providing information about the biological effects of perturbations. However, when a perturbation affects multiple factors, it can be difficult to synthesize the effects of this perturbation across all affected factors. GSFA provides a way to integrate information over all factors to compute the total effect of a target perturbation on individual genes. Under our model, this total effect is simply the product of the perturbation-to-factor effect and the gene-on-factor loading, summing over all factors (Fig. 1). The significance of the summarized total effect is evaluated using local false sign rate (LFSR)^26^, a simple summary of the posterior distribution with an interpretation similar to local false discovery rate (Methods).

In applying GSFA to scRNA-seq data, we first converted the raw UMI counts into deviance residuals^27^, a continuous quantity analogous to z-scores that approximately follows a normal distribution. The deviance residual transformation overcomes some problems of the commonly used log transformation of read counts, and has been shown to improve downstream analyses, such as feature selection and clustering (Supplementary Note). GSFA produces three main outputs (Fig. 1, bottom): the association between genetic perturbations and factors; the weights of genes on factors measured by PIPs; and a list of DEGs of each perturbation at a given LFSR cutoff. In cases where the experiment involves multiple cell types or conditions, GSFA can estimate these effects separately and produce different DEGs for each cell type/condition (Supplementary Note).

### Simulation study demonstrates the advantages of GSFA

We performed extensive simulations to evaluate the performance of GSFA under two settings. In the first simulation setting, referred to as the “normal distribution scenario”, we generated continuous gene expression levels with a normal error distribution, according to the GSFA model (Methods). Each dataset consists of 4000 cells, 6000 genes, 6 types of perturbations, and 10 underlying factors. Each perturbation occurs in ∼5% of cells, mimicking real multiplex CRISPR screening assays. The proportion of genes with non-zero effects on each factor, referred to as factor density below, varies from 5% to 20%. For simplicity, each perturbation is associated with a distinct factor. The second “count-based” simulation setting mimics real scRNA-seq UMI data. We converted normally distributed expression levels into count data following a Poisson model incorporated with varying library sizes (Methods). Other simulation parameters remained the same.

Simulated data allowed us to evaluate the model choice, particularly the prior distribution on gene weights (W) in count-based data. From our simulations, factors inferred under the spike-and-slab prior sometimes resulted in factors much denser than the ground truth. In contrast, factors inferred under the normal-mixture prior have sparse gene weights, with the proportion of loaded genes centered around the ground truth (Fig. S1a). This justifies our choice of normal-mixture prior as the default prior for read count data.

To evaluate the performance of GSFA in the inference of factors, we quantified the correlation between the inferred factors with true factors. Across all scenarios, factors inferred by GSFA are highly correlated with the true underlying factors, with 80-90% of the absolute correlation values above 0.8 (Fig. 2a,b). GSFA also recovers genes with nonzero loading on the factors. Indeed, genes with PIPs above 0.95 are generally true genes, with observed false discovery proportions (FDP) below 0.1 when the true factor density is less than 0.2 (Fig. S1b,c).

**Figure 2:**
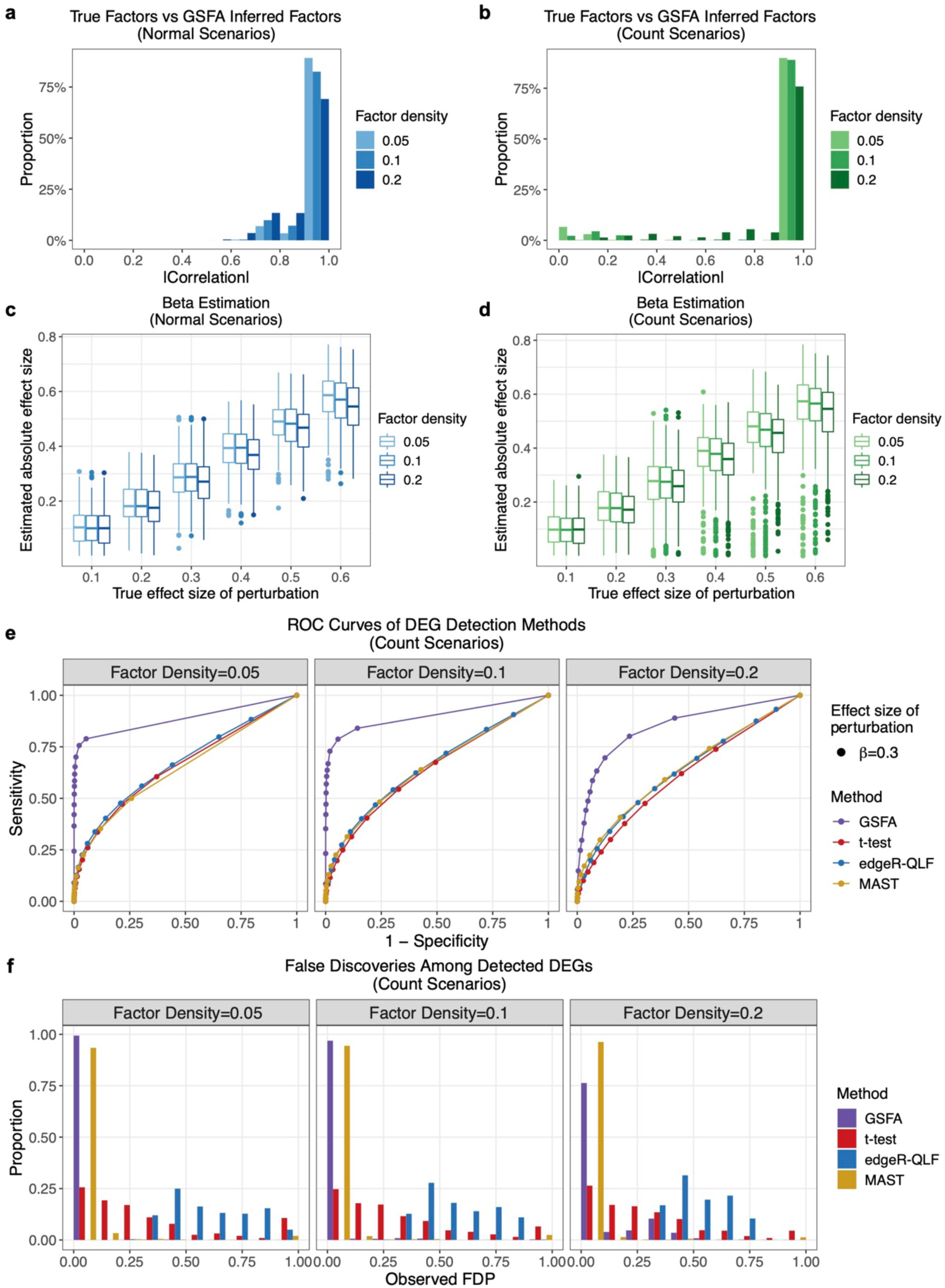
GSFA performance on simulated data. **a**) Distributions of the absolute correlation values between true factors and the factors inferred by GSFA under the normal setting. Different colors represent different values of true factor density varying from 0.05 to 0.2. **b**) Same as in a) but under count-based scenarios. **c**) Box plots of absolute effect sizes from perturbation-factor regression estimated by GSFA under the normal setting. Different colors represent different values of true factor density varying from 0.05 to 0.2. **d**) Same as in c) but under count-based scenarios. **e**) ROC curves of DEG discovery under the count-based setting and 3 different levels of true factor density; 4 colors correspond to 4 DEG detection methods. Results shown are of perturbations with a true association effect of 0.3 on factors. See Fig. S2 and S3 for results under other settings. **f**) Distributions of observed proportion of false discoveries (FDP) among significant DEGs detected by GSFA (LFSR < 0.05) and other methods (FDR < 0.05) per dataset under the count-based setting and various true factor densities. 4 colors correspond to 4 DEG detection methods.

Next, we evaluated the performance of GSFA in detecting the effects of perturbations on factors. Across all scenarios, GSFA estimates these effects accurately (Fig. 2c,d). The small downward bias of estimated effects is expected, given the sparse prior we imposed. We further assessed the calibration of PIPs of these effects. At a PIP threshold of 0.95 and a true factor density level below 0.2, the proportion of falsely detected effects is generally below 0.1 (Fig. S1d,e). Finally, we compared the performance of GSFA in DEG detection, *i*.*e*., detection of genes affected by perturbations, with commonly used differential expression analysis methods: the Welch’s t-test^28^, the edgeR quasi-likelihood F-test (edgeR-QLF)^13^, and MAST, a method designed for single cell analysis^29^. We generated receiver operating characteristic (ROC) curves for each method by varying the LFSR or false discovery rate (FDR) threshold. GSFA outperformed the other methods in terms of both sensitivity and specificity under all scenarios (Fig. 2e, Fig. S2, Fig. S3). In addition, DEGs detected by GSFA at LFSR < 0.05 have observed FDPs well below 0.05 in most cases, while the observed FDPs among edgeR or t-test DEGs show significant inflation under the count-based scenarios (Fig. 2f).

Through these simulations, we have demonstrated that GSFA is a powerful method to identify gene modules, the associations between perturbations and modules, and the specific genes affected by each perturbation.

### GSFA reveals downstream effects of regulators of T cell activation

We applied GSFA to a CROP-seq dataset of primary human CD8^+^ T cells^10^. Using a series of CRISPR screens, the authors identified a dozen genes that regulate T cell proliferation in response to T cell receptor (TCR) stimulation. They then used CROP-seq to target these genes, along with 8 known immune checkpoint genes, in stimulated and unstimulated T cells, and applied a clustering approach to characterize the effects of each perturbation. Although the authors found that perturbations of some genes were correlated with clusters characterized by T cell activation or resting states, many other genes were not associated with any cluster. Moreover, the study lacked systematic differential expression analysis to reveal specific genes affected by perturbations.

To fill in these gaps, we applied GSFA to this CROP-seq dataset, allowing perturbations to have different effects on factors in stimulated and unstimulated cells (Methods). With a total of 20 factors pre-specified in GSFA, we obtained 24 associations (PIP > 0.95) between perturbations and factors in stimulated cells that involved 9 gRNA-targeted genes (Fig. 3a for a subset of factors, full results in Fig. S4a). Among these genes, the effects of *ARID1A, DGKA, SOCS1*, and *TCEB2* were undetected by clustering analysis in the original study (Fig. 3b). In unstimulated cells, only three pairs of associations were detected at PIP > 0.95 (Fig. S4b), which is unsurprising given the role of these targeted genes in regulating T cell responses. Nevertheless, it is interesting that some perturbations (*e*.*g. TCEB2, RASA2*) have effects in unstimulated cells (Fig. S4b), suggesting that these genes affect resting state transcriptomes. We also confirmed, with permutation analysis, that the full GSFA results, including the inferred perturbation effects and gene loading, were calibrated (Fig. S5a-d). Altogether, these results highlight the power of GSFA to detect the broad effects of target genes on the latent factors.

**Figure 3:**
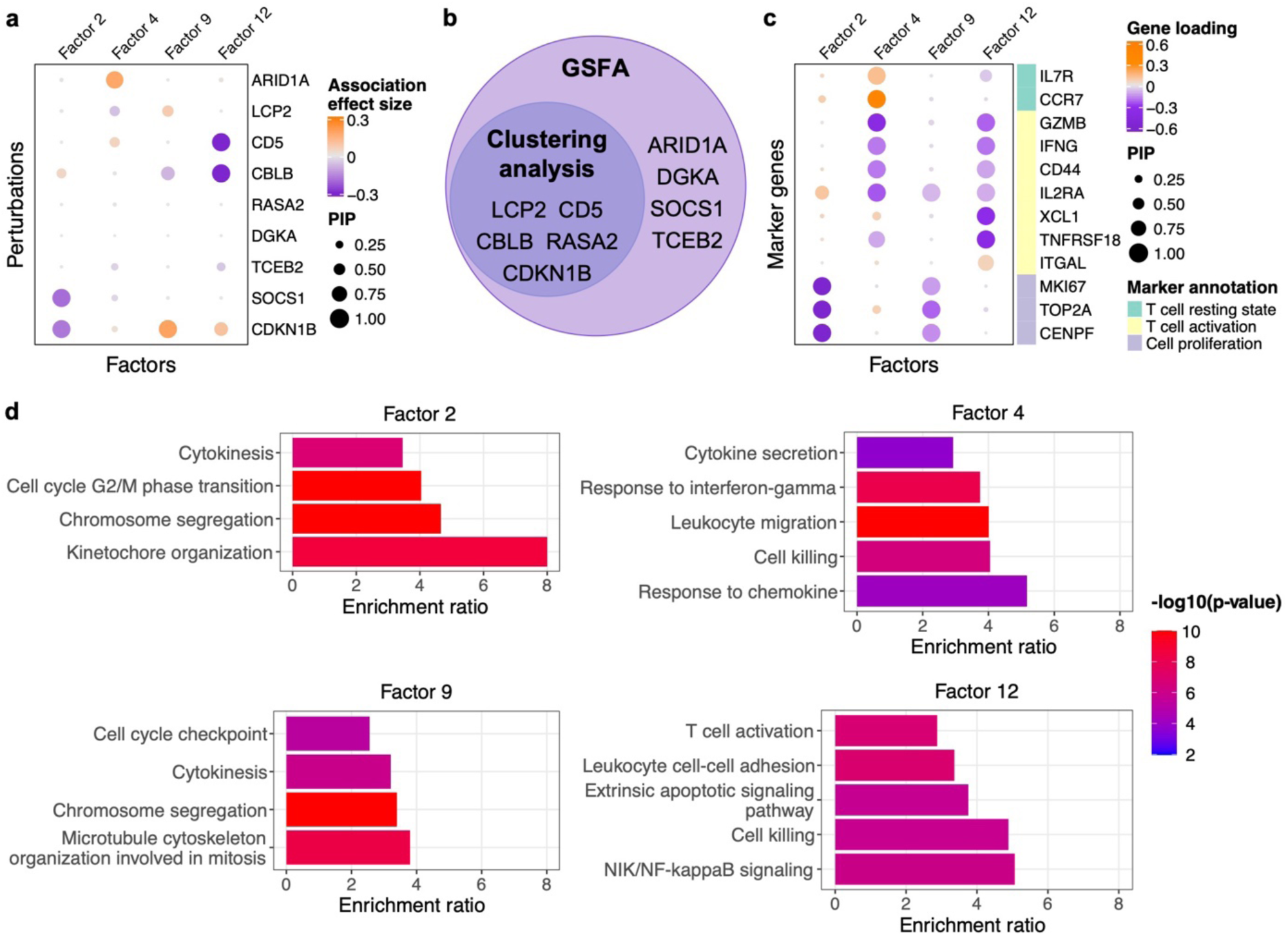
GSFA results of inferred factors from analysis of CROP-seq data of primary CD8^+^ T cells. The results were based on stimulated T cells. **a**) Estimated effects of gene perturbations on selected factors inferred by GSFA. The size of a dot represents the PIP of association; the color represents the effect size. **b**) Venn diagram of targets identified using the original clustering-based method vs. GSFA. **c**) Loading of selected marker genes on selected factors. The size of a dot represents the gene PIP in a factor and the color represents the gene weight (magnitude of contribution) in a factor. **d**) The fold of enrichment for selected GO “biological process” gene sets significantly enriched (q-value < 0.05) in factor 2, 4, 9, and 12. Bars are colored by −log_10_ p-values from the over-representation tests, where genes with PIP > 0.95 in the factor were compared against a background of all genes used in GSFA.

To characterize these latent factors – particularly those associated with perturbations – we inspected the weights of canonical marker genes and performed GO enrichment analysis of genes loaded on the factors. As an example, factors 2 and 9 have negative weights for cell proliferation markers *MKI67, TOP2A*, and *CENPF* (Fig. 3c), and are enriched for GO terms related to cell cycle and division (Fig. 3d). Factors 4 and 12 are associated with markers of T cell activation and/or resting states (Fig. 3c) and are enriched for GO terms related to immune responses (Fig. 3d). Together, these results show that the latent factors discovered by GSFA represent cellular processes.

We note that a target gene may affect multiple factors representing related processes. For instance, targeting of *CDKN1B* is associated with two cell-cycle-related factors with opposite signs (Factors 2 and 9; Fig. 3a,c). This makes it difficult to understand target genes’ effects. We thus used GSFA’s LFSR-based differential expression analysis (Fig. 1) to identify specific downstream genes affected by the perturbations. We also ran several other differential expression analysis methods for comparison, including scMAGeCK-LR^30^, a method tailored for single-cell CRISPR screening data, MAST^29^, DESeq2^12^, and edgeR-QLF^13^. scMAGeCK-LR and MAST had calibrated false positive rates in the permuted data, while DEseq2 showed modest inflation and edgeR-QLF severe inflation (Methods, Fig. S5e-h). Therefore, we excluded edgeR-QLF from further analysis. In stimulated T cells, GSFA detected 148 to 710 DEGs at LFSR < 0.05 for 9 gene targets, four of which (*ARID1A, DGKA, SOCS1*, and *TCEB2*) were poorly characterized by clustering analysis in the original study^10^. Compared with other methods, GSFA consistently detected the most DEGs across these 9 targets, sometimes with 10 times or more DEGs (Fig. 4a). Additionally, the DEGs of all 9 target genes detected by GSFA are enriched for a larger number of GO terms (Fig. 4b), many of which are relevant to T cell responses (Fig. 4c). DEGs detected by other methods, in contrast, have almost no GO enrichment except for *TCEB2*. These results thus show that GSFA has much higher sensitivity in detecting DEGs than existing methods.

**Figure 4:**
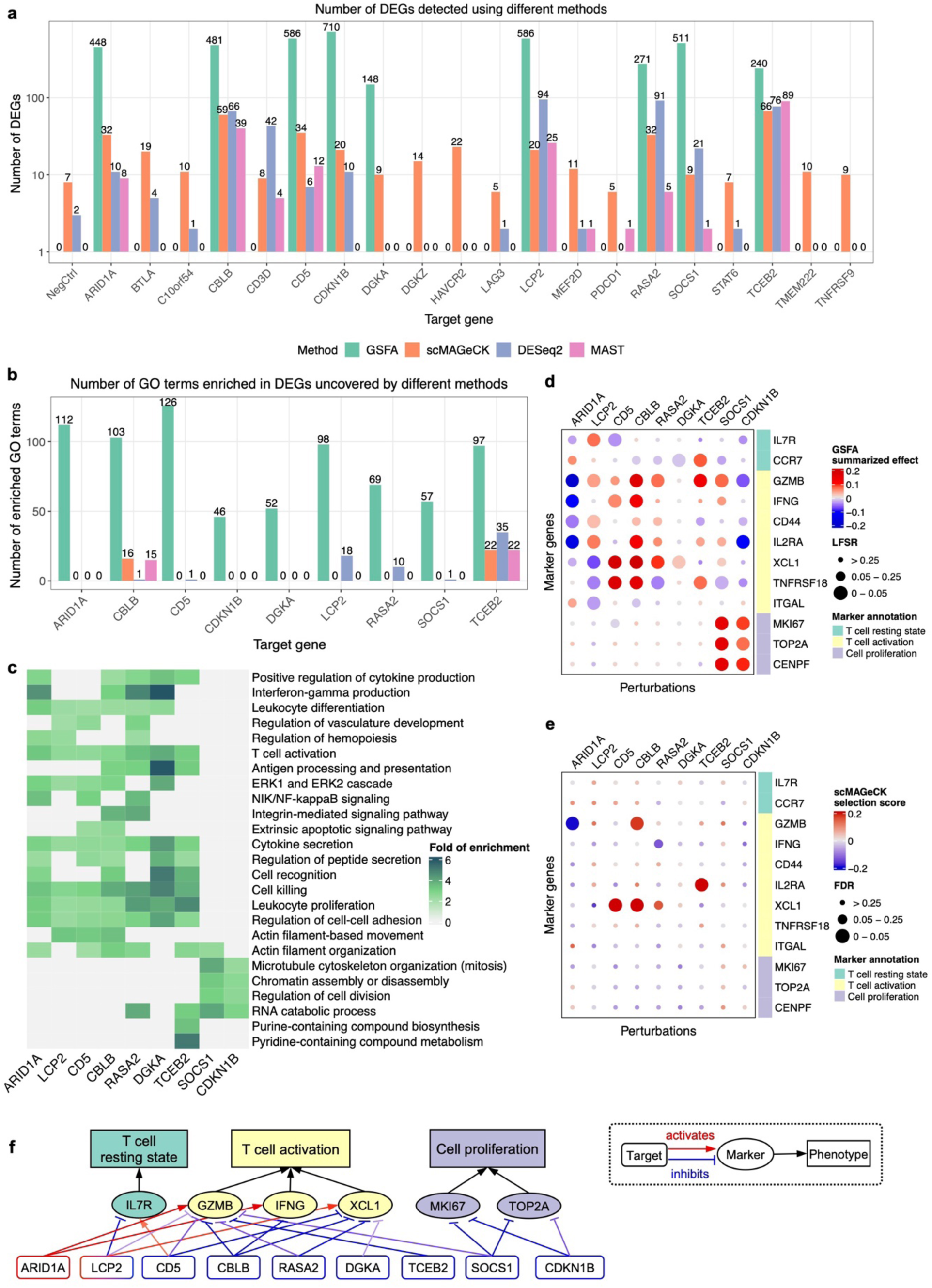
GSFA results of the effects of genetic perturbations on gene expression in CD8^+^ T cell data. The results were based on stimulated CD8^+^ T cells. **a**) Number of DEGs detected under all perturbations using 4 different methods. The y axis is log scaled and bar height corresponds to count+1 (as the number of DEGs could be 0); the exact number of DEGs are labeled on top of the bars. The detection threshold for DEGs is LFSR < 0.05 for GSFA, and FDR < 0.05 for all other methods. **b**) Number of GO Slim “biological process” terms enriched in DEGs detected by different methods. **c**) Heatmap of selected GO “biological process” terms and their folds of enrichment in DEGs (LFSR < 0.05) detected by GSFA under different perturbations. **d**) GSFA estimated effects of perturbations on marker genes in stimulated T cells. Sizes of the dots represent LFSR bins; colors of the dots represent the summarized effect sizes. **e**) scMAGeCK estimated effects of perturbations on marker genes in stimulated T cells. Sizes of the dots represent FDR bins; colors of the dots represent the scMAGeCK selection scores. **f**) A target-marker-phenotype regulatory network summarizing GSFA results. Significant (LFSR < 0.05) regulatory relationships between target and marker genes are represented by colored arrows, with red sharp arrows indicating positive regulation of marker genes by the target genes, and blue blunt arrows indicating negative regulation. The darkness of color represents the relative magnitude of effect. Note the effect directions here are the opposite of the perturbation effects.

With a large number of DEGs detected for the 9 target genes, we next focused on their effects on specific marker genes of T cell activation to better understand the genes’ functions. GSFA revealed a number of effects of the target genes on the markers (Fig. 4d), many of which were missed by other methods (Fig. 4e for scMAGeCK; Fig. S4d,e for others). The estimated effects of these genes on the chosen markers largely agree with the reported roles of these genes^10^. For instance, targeting of *CD5, CBLB* and *RASA2* have mostly positive effects on markers of activated T cells, and negative or no effects on markers of resting T cells (Fig. 4d), consistent with the functions of these genes as negative regulators of T cell stimulation^10^. Targeting of *CDKN1B* has strong positive effects on cell proliferation markers (Fig. 4d), consistent with its function in the cell cycle^31^ and its role as a negative regulator of T cell proliferation^10^.

Our analysis also revealed molecular effects of the four novel genes, *ARID1A, DGKA, SOCS1*, and *TCEB2*, whose effects were poorly characterized in the earlier study (Fig. 3b). The effects of perturbations of *TCEB2* and *DGKA* on T cell markers are similar to those of other negative regulators of T cell responses, such as *CD5*. Targeting *SOCS1* has a strong effect on cell proliferation markers (Fig. 4d). Accordingly, several genes of the SOCS (suppressor of cytokine signaling) family have been reported to inhibit cell cycle progression^32^. Targeting of *ARID1A*, a chromatin remodeler and potential tumor suppressor^33–35^, has strong negative effects on the effector markers (Fig. 4d), suggesting its role as a positive regulator of T cell activation. Indeed, *ARID1A* mutations occur in many human cancer types, and result in limited chromatin accessibility and down-regulation of interferon-responsive genes, leading to poor tumor immunity^36^.

Collectively, GSFA revealed detailed transcriptional effects of 9 genetic perturbations, including four genes largely missed by clustering or differential expression analysis with other tools. The functions of these four genes suggested by GSFA analysis are largely consistent with their known biological roles. We constructed a regulatory network to summarize our major findings of the functions of all nine target genes (Fig. 4f). Our results highlight the power of GSFA in revealing the detailed molecular effects of genetic perturbations in single cell CRISPR screens.

### GSFA characterizes the transcriptomic effects of autism risk genes

We next applied GSFA to CROP-seq data targeting 14 neurodevelopmental genes, including 13 autism risk genes, in LUHMES human neural progenitor cells^37^. After CRISPR targeting, the cells were differentiated into postmitotic neurons and sequenced. The authors then projected cells onto a pseudotime trajectory, which approximates the progression of neuronal differentiation, and associated the perturbations with the pseudotime of cells. This analysis revealed the effects of several target genes on neuronal differentiation. However, it provided limited information on the molecular processes affected by the target genes other than pseudotime, and its differential expression analysis did not take measures to control the false discovery rate.

After applying GSFA to this dataset, we first confirmed that the GSFA results were calibrated, *i*.*e*., GSFA did not produce false positive findings in permutations (Fig. S7). We found significant effects (PIP > 0.95) of 7 target genes, including *ADNP, ARID1B, ASH1L, CHD2, DYRK1A, PTEN*, and *SETD5*, on at least one out of 20 latent factors (Fig. 5a for a subset of factors, Fig. S6a for full results). Among the 7 genes, the transcriptomic effects of *ADNP* and *SETD5* were missed in the original pseudotime-based analysis (Fig. 5b). We characterized these factors by inspecting the weights of neuronal markers and GO enrichment analysis. In factor 4, for example, the markers of mature neurons such as *MAP2* and *NEFL* have negative weights, while negative regulators of neuron projection such as *ITM2C* have positive weights (Fig. 5c), suggesting that factor 4 is negatively associated with neuronal maturation. Indeed, factor 4 is significantly enriched for gene sets involved in neuronal development (Fig. 5d). Factors 9 and 16, similarly, show loadings of neuronal markers and are enriched for relevant GO terms (Fig. 5c,d). These results suggest that GSFA was able to relate genetic perturbations with biologically meaningful latent factors.

**Figure 5:**
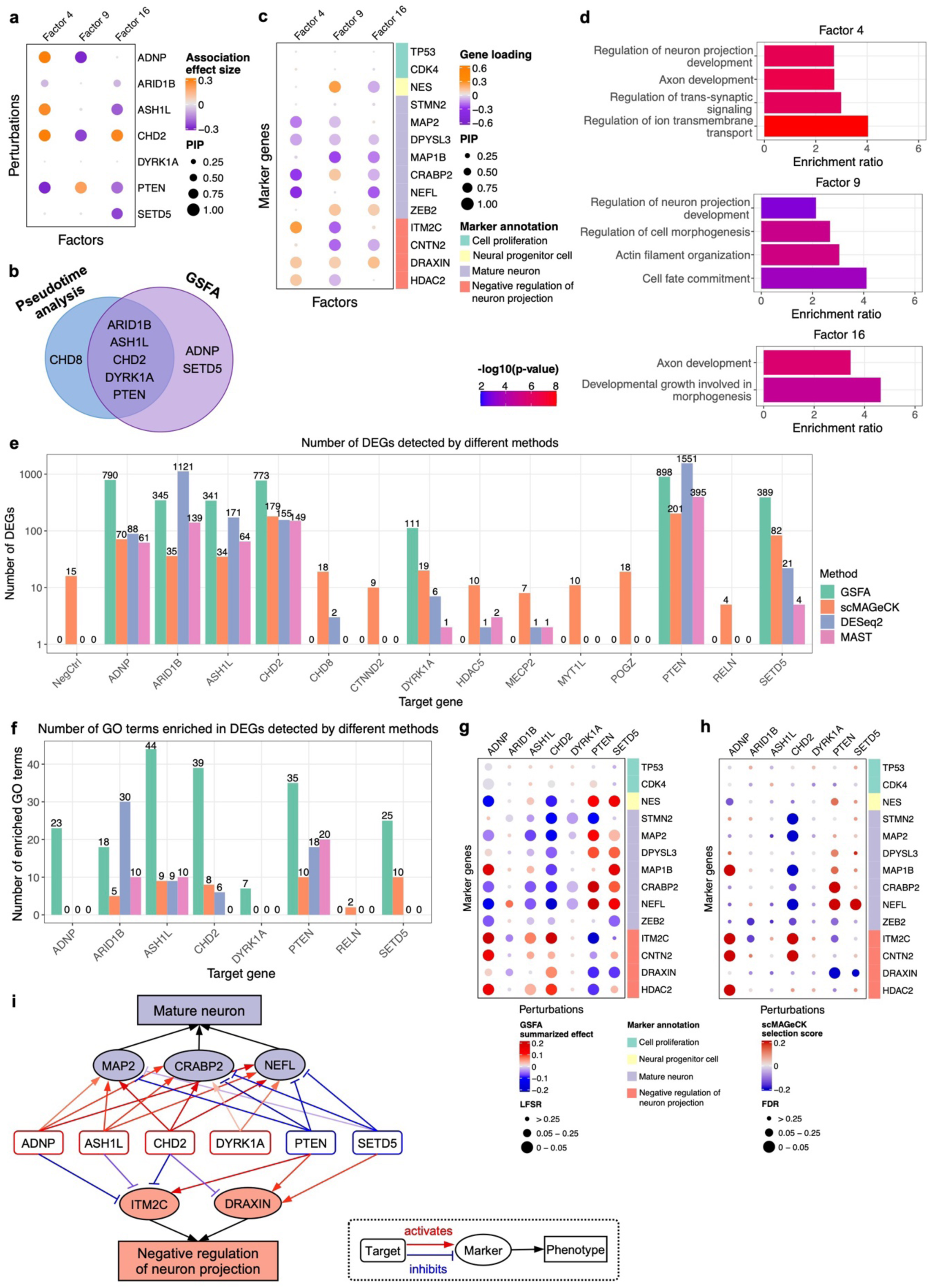
GSFA analysis of the CROP-seq data of LUHMES cells. **a**) Estimated effects of gene perturbations on selected factors inferred by GSFA. The size of a dot represents the PIP of association; the color represents the effect size. **b**) Venn diagram of targets identified from the original pseudotime association analysis vs. from GSFA. **c**) Loading of neuronal marker genes on selected factors. The size of a dot represents the gene PIP in a factor and the color represents the gene weight (magnitude of contribution) in a factor. **d**) The fold of enrichment for selected GO “biological process” terms enriched in factor 4, 9, and 16 (q-value < 0.05). Bars are colored by −log_10_ p-values from the over-representation tests, where genes with PIP > 0.95 in the factor were compared against a background of all genes used in GSFA. **e**) Number of DEGs detected under all perturbations using 4 different methods. The y axis is log scaled and bar height corresponds to count+1 (as the number of DEGs could be 0); the exact number of DEGs are labeled on top of the bars. The detection threshold for DEGs is LFSR < 0.05 for GSFA, and FDR < 0.05 for all other methods. **f**) Number of GO Slim “biological process” terms enriched in DEGs detected by different methods. **g**) GSFA estimated effects of perturbations on marker genes. Sizes of the dots represent LFSR bins; colors of the dots represent the summarized effect sizes. **h**) scMAGeCK estimated effects of perturbations on marker genes. Sizes of the dots represent FDR bins; colors of the dots represent the scMAGeCK selection scores. **i**) A target-marker-phenotype regulatory network summarizing GSFA results. Significant (LFSR < 0.05) regulatory relationships between target and marker genes are represented by colored arrows, with red sharp arrows indicating positive regulation of marker genes by the target genes, and blue blunt arrows indicating negative regulation. The darkness of color represents the relative magnitude of effect. Note that the direction of regulation is the opposite of the perturbation effect.

We next identified the individual genes affected by the perturbations. GSFA detected DEGs at LFSR < 0.05 for the same 7 gene targets (Fig. 5e). Compared with other differential expression analysis methods, GSFA detected the most DEGs for 5 out of 7 gene targets (Fig. 5e). Furthermore, DEGs detected by GSFA are enriched for the most GO terms across almost all targets (Fig. 5f), many of which are related to neuronal development or neural signaling (Fig. S6e).

To better understand the functions of these 7 target genes, we examined their effects on marker genes for neuron maturation and differentiation. GSFA uncovered perturbation effects on a number of neuronal marker genes across all targets except *ARID1B* (Fig. 5g), while other methods detected fewer differentially expressed markers (Fig. 5h for scMAGeCK, Fig. S6c,d for MAST and DESeq2). GSFA-estimated effects of the target genes largely validated the known functions of these genes. Targeting of *ASH1L, CHD2*, and *DYRK1A* has mostly negative effects on mature neuronal markers, and positive effects on negative regulators of neuron projection (Fig. 5g), indicating delayed neuron maturation by the repression of these genes. Knockdown of *PTEN* showed opposite effects by GSFA, suggesting its opposite role on neuronal differentiation. These results agree with the experimental findings of the effects of these perturbations on neuronal maturation phenotypes^37^.

Two genes, *ADNP* and *SETD5*, were missed in the pseudotime-based analysis in the original study (Fig. 5b). The estimated effects of these genes on neuronal markers by GSFA suggested that repression of *ADNP* would lead to delayed neuronal differentiation, whereas *SETD5* repression would have the opposite effect (Fig. 5g). These predictions are consistent with the experimental finding of *ADNP* in the original study^37^, and with the finding that *SETD5* knockdown increases the proliferation of cortical progenitor cells and neural stem cells^38^.

In conclusion, GSFA allowed us to identify and characterize the transcriptional effects of 7 ASD risk genes, including *ADNP* and *SETD5*, whose effects were largely missed in the original study. Detailed characterization of these genes by GSFA using neuronal markers provides functional insight that is largely consistent with their known roles in literature. While GSFA missed the effect of *CHD8* (Fig. 5b), we noticed that all the existing DEG methods largely missed its effect as well (Fig. 5e). We summarized the inferred target effects by GSFA on selected marker genes and affected cellular processes in a gene regulatory network (Fig. 5i). Together, these results confirm the advantages of GSFA in analyzing single cell screening data.

## Discussion

Single-cell CRISPR screening technologies have enabled efficient readouts of transcriptome-level effects of multiple genetic perturbations in tens of thousands of individual cells in a single experiment. These technologies offer great opportunities, but also challenges for effective data analysis. We presented GSFA to address these challenges. GSFA can identify biologically interpretable gene modules that respond to genetic perturbations, and by summarizing the information from all factors, GSFA can infer the effects of perturbations on specific downstream genes. When applied to two CROP-seq datasets, GSFA detected transcriptomic effects and shed light on the molecular mechanisms of regulators of T cell activation and neuronal differentiation, respectively.

GSFA is built on factor analysis, which we believe offers key benefits over clustering-based analysis of single-cell screening data^39,40^. Clustering-based analysis requires the effects of perturbations to be large enough to alter cluster compositions; thus, it may miss perturbations with moderate effects. In contrast, factor analysis does not rely on disjoint cell clusters and can potentially detect subtler effects, *e*.*g*. those of non-coding regulatory elements. In addition, as we have demonstrated, the inferred factors lead to better biological interpretability than cell clusters. Conceptually, a researcher can perform factor analysis or related methods such as topic modeling on the expression data, and then correlate the inferred factors with genetic perturbations across cells. This approach was used in the MUSIC method^41^. Compared with this two-step approach, GSFA has several advantages. When inferring expression factors, GSFA uses the genetic perturbation as a prior to improve the estimation of the factors (hence “guided” in the method name, see Methods, also discussion below). In practice, GSFA offers an important advantage when a perturbation affects multiple factors. For example, in the LUHMES data, target genes often show associations with two or more factors (Fig. 5a, S6a), which may have overlapping functions (Fig. 5d). Thus, a perturbation may have positive effects on some factors, and negative effects on others. Similarly, some factors may correlate positively with the biological process they represent, *e*.*g*. cell maturation, while others correlate negatively. In such cases, it would be extremely difficult to learn the total effects of the perturbation. GSFA solves this problem by synthesizing the effects of a perturbation over all factors.

In GSFA, factors are dependent on the genetic perturbations via a linear model. In this sense, GSFA is related to a class of factor models in the statistics literature, sometimes called supervised factor analysis, where the factors depend on covariates of samples^42–44^. These models can help improve the estimation of latent factors, and have been proposed in bulk gene expression data analysis^45^ where samples have different characteristics or experimental conditions. Nevertheless, existing covariate-dependent factor models were designed only for factor inference, and do not provide estimates of the effects of covariates (perturbations in our case) for specific genes, *i*.*e*., they cannot estimate perturbation effects for single-cell CRISPR screening data.

GSFA is a versatile tool for single-cell screening data analysis and can be potentially used in other settings. As demonstrated in our study of CD8^+^ T cell data, GSFA can be used in datasets with multiple cell types or treatment conditions, and estimate the perturbation effects for each cell group separately. Secondly, while both applications used the low multiplicity of infection (MOI) setting, with each cell typically harboring at most one gRNA perturbation, GSFA’s linear model of how factors depend on genetic perturbations easily accommodates multiple perturbations per cell, and thus can be readily applied to high MOI settings. Thirdly, the implementation of GSFA in Rcpp makes it relatively efficient (Methods). With ∼10k cells in our current analysis, GSFA takes several hours. Finally, the generality of the statistical model of GSFA makes it applicable to bulk RNA-seq studies with paired genetic perturbations, *e*.*g*., cancer transcriptomic data with somatic mutations, or expression quantitative trait locus (eQTL) data with paired genotype-expression measurements.

GSFA can be further improved along several directions. GSFA does not directly model read counts and instead uses deviance residuals converted from count data. While this transformation generally works well in our analysis, we noticed that LFSRs from differential expression analysis can be modestly inflated at high factor density (Fig. 2f, under *π* = 0.2). Hence, directly modeling the read count data may improve the calibration and power of GSFA. Another limitation of GSFA is that we assume genetic perturbations affect downstream genes through factors. It is possible that the factors may not fully capture the transcriptional effects; thus, it may be desirable to add to the model “direct effect’’ terms, where perturbations directly affect the expression of a gene without acting on any factors. Finally, GSFA uses Gibbs sampling for inference; replacing this with a more efficient algorithm, such as variational approximation, may reduce the computational time.

In conclusion, single cell CRISPR-screening is a promising technology, yet the difficulty of data analysis has prevented us from realizing its full potential. GSFA complements the strength of this technology by offering a powerful new analysis framework, representing a substantial advance of the field.

## Methods

### GSFA model

The input data of GSFA consist of a gene expression matrix **Y**_*N*×*P*_ with *N* cells and *P* genes, and a perturbation matrix **G**_*N×P*_ with *N* cells and *M* types of genetic perturbations. In all our analyses, the perturbation matrix is binary, *i*.*e*., *G*_*ij*_ =1 if cell *i* has the *m*-th type of perturbation and 0 otherwise, but this is not strictly required by the model, *e*.*g*., **G** might represent the dosage of genetic perturbations. The GSFA model has two main parts: (1) a sparse factor analysis model that decomposes the expression matrix **Y** into a factor matrix **Z**_*N×K*_, where *K* is the number of factors, and a sparse gene weight matrix **W**_*P*×*K*_, and (2) a multivariate linear model that correlates the factor matrix **Z** with the perturbation matrix **G**:

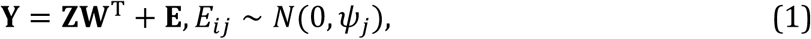

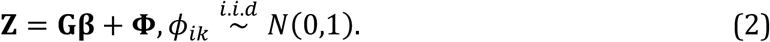

Here **E** is an *N* × *P* residual matrix with gene-specific variances stored in a *P*-vector ***ψ*, β** is an *M* × *K* matrix of perturbation effects on factors, and **Φ** is an *N* × *M* residual matrix with variance 1. Compared with standard factor-analysis, our model assumes the latent factors **Z** also depend on additional covariates **G**, hence our model is a form of “guided” factor analysis.

We assume that each perturbation affects only a small number of factors, so we impose a “spike-and-slab” prior on the effect of perturbation *m* (1 ≤ *m* ≤ *M*) on factor *k* (1 ≤ *k* ≤ *K*):

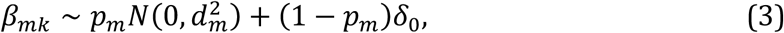

where *δ*_0_ is delta-function, *p*_*m*_ denotes the proportion of factors affected by perturbation *m*, and *d*_*m*_ the prior variance of the effect sizes of *m*.

To limit the number of genes contributing to a factor and facilitate the biological interpretation of factors, we also impose a sparse prior on the gene weights. We found in our simulations and real data analysis that, when analyzing count data, the standard spike-and-slab prior is sometimes insufficient to impose sparsity, so we chose a “normal-mixture” prior (see the main text). This prior assumes that the gene weights in a factor come from a mixture of two normal distributions with mean 0 but different variances. The difference with the spike-and-slab prior is that the “background” component is not necessarily *δ*_0_, but rather a distribution with small effects. The prior weight of gene *j* in the factor *k* follows:

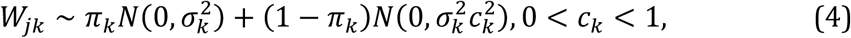

where *π*_*k*_ represents the proportion of genes affected by the factor *k* (the “foreground” part), and 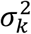 the prior effect size variance of factor *k*, and *c*_*k*_ a scale parameter controlling the relative size of the foreground and background effects.

The prior distributions for other parameters in the model are specified in Supplementary Note 1.1.

### GSFA model inference

We infer the parameters in GSFA using Gibbs sampling, a Markov chain Monte Carlo (MCMC) algorithm that obtains a sequence of approximate samples from their posterior distribution given the observed data. Gibbs sampling is an attractive choice here because the conditional distributions of the main parameters (**β** and **W**) and latent variables (**Z**) have analytic forms. To see this, we first consider the conditional distribution of **W**, given data and all other parameters/variables, P(**W**|**Y**,**G**,**Z**,**β**) (for simplicity, we drop the hyperparameters and parameters related to the error terms). It’s easy to see that given **Z, W** does not depend on **G** and **β**, so we have:

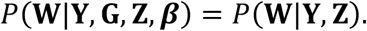

The problem now becomes multivariate linear regression, **Y = ZW**^**T**^ **+ E**, where **W** follows a spike-and-slab prior. This is a well-studied problem in the statistics literature^46,47^. Similarly, we can see that the conditional distribution of **β** is given by:

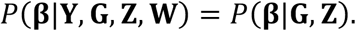

Again, this reduces to a regression problem **Z = Gβ + Φ**, where **β** follows the normal-mixture prior. Finally, the conditional distribution of **Z** is given by:

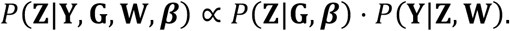

This is also a regression problem **Y** = **ZW**^T^ + **E**, where **Z** represents the unknown coefficients, with a normal prior, *Z*_*i*_ ∼ *N*(*G*_*i*_**β, I**), for the sample *i* (1 ≤ *i* ≤ *N*). We now see that the posterior of Z not only depends on gene expression matrix **Y**, but also the perturbations **G**. In other words, the perturbations impose a prior on **Z**, hence “guiding” the inference of **Z** in a certain sense.

To facilitate computation, we also introduced two latent binary matrices, **F**_*P* × *K*_ and **γ**_*M* × *K*_, to indicate which distribution the corresponding parameters in **W** and **β** come from. The joint prior distribution of **W** and **F** follows:

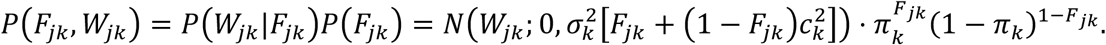

The joint prior distribution of **β** and **γ** can be written as:

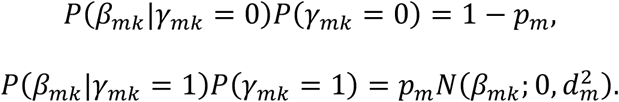

The details of the Gibbs sampling steps are described in Supplementary Note 1.2.

Unless mentioned otherwise, for all the datasets in the study, we ran the MCMC chain for 3000 iterations, and used the last 1000 iterations to obtain the posterior samples of parameters.

The posterior distribution allows us to summarize the probabilities that some effects are non-zero. Specifically, the posterior mean of *γ*_*mk*_ gives the posterior inclusion probability (PIP) of *β*_*mk*_, *i*.*e*., the probability of *β*_*mk*_ being non-zero as:

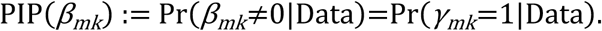

Similarly, the posterior mean of *F*_*jk*_ gives the PIP of *W*_*jk*_ defined as the probability of *W*_*jk*_ coming from the “foreground” normal distribution given data:

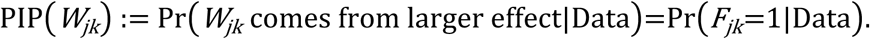

### Summarizing the effects of genetic perturbations on individual genes

While the effects of genetic perturbations are formulated in terms of factors under GSFA, the model does allow us to infer the effects on individual genes. This is similar to the commonly used differential gene expression analysis, where the expression of genes in cells with certain perturbation are compared with those without it. Under our model, the effect of perturbation *m* on the expression of gene *j* is mediated through one or more factors. The total effect, denoted as *θ*_*mj*_, is then given by the sum of *K* mediated effects:

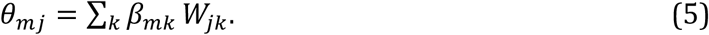

To sample the posterior distribution of *θ*_*mj*_, we use the posterior samples of *β*_*mj*_ and *W*_*jk*_:

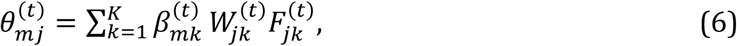

where superscript (*t*) denotes the *t*-th posterior sample. While the posterior distribution of *θ*_*mj*_ contains all the information we have, in practice, it is simpler to use a single summary of how likely *θ*_*mj*_ is non-zero. To do this, we use the local false sign rate (LFSR), a metric that is analogous to local false discovery rate (LFDR), but reflects confidence in the sign of effect rather than in the effect being non-zero^26^. It has been shown that LSFR has some benefits over the commonly used FDR approach, and is in fact more conservative than LFDR. The LFSR of the perturbation effect on individual genes, *θ*_*mj*_, is given by:

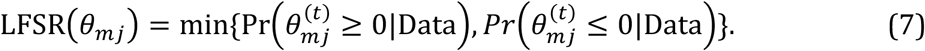

By thresholding LFSR, we can obtain significant DEGs under each perturbation. In practice, the threshold is LFSR < 0.05.

### GSFA implementation and running time

The computational complexity of the GSFA inference is *O*(*NP*) per iteration, with *N* being the number of cells and *P* being the number of genes. The average run time of GSFA on a simulated dataset of 4000 cells and 6000 genes is 1.32 seconds per iteration on a modern Linux workstation with Intel Xeon E5-2680 v4 (2.40 GHz) processors. GSFA was implemented in R and Rcpp. The R package is available at Github: https://github.com/gradonion/GSFA.

All analyses in this study were performed in `R 4.0.4`.

### Alternative differential gene expression (DGE) methods

For comparison, we applied the following DGE methods to simulated or real data:

1. Welch’s t-test^28^ using the `t.test()` function in R;
2. edgeR quasi-likelihood F-test (edgeR-QLF)^13^ using `glmQLFit()` and `glmQLFTest()` functions in the R package `edgeR`;
3. DESeq2^12^ using the `DESeq()` function in the R package `DESeq2`;
4. MAST^29^, a statistical method tailored for scRNA-seq data, using `zlm()` and `lrTest()` functions in the R package `MAST`;
5. scMAGeCK-LR^48^, a linear-regression-based approach tailored for single-cell CRISPR screening data, using the `scmageck_lr()` function in the R package `scMAGeCK`. We did not include scMAGeCK-RRA as it is not designed to test all genes^48^.

### Simulation study

We simulated single-cell CRISPR screen data under several scenarios, with either continuous gene expression levels or discrete gene count data as output. We simulated under *N* = 4000 cells, *P* = 6000 genes, *M* = 6 types of perturbations, and *K* = 10 underlying factors.

1. Normal model Continuous gene expression levels were generated under the following model:

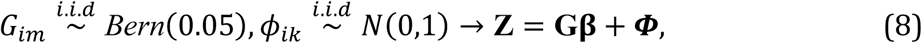

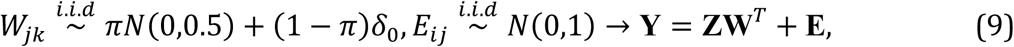

where *π* represents the proportion of genes loaded on any factor, and varies from 0.05, 0.1 to 0.2 under different simulation scenarios.
2. Count model To sample the read count data, we assume each cell has a library size/scaling factor *L*_*i*_, sampled from a normal distribution with mean 5×10^5^. The count of a gene *j* would then be sampled from a Poisson distribution with its mean determined by the continuous gene expression level *y*_*ij*_ and the scaling factor *L*_*i*_:

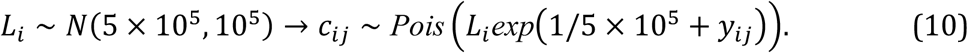

The sampled counts are converted to deviance residuals (Supplementary Note 2.1), then centered and scaled so that each gene has variance 1 before provided as input for GSFA.

We set the effect size matrix **β** to the following form so that each perturbation affects a distinct factor, and the effect sizes vary from 0.1 to 0.6.

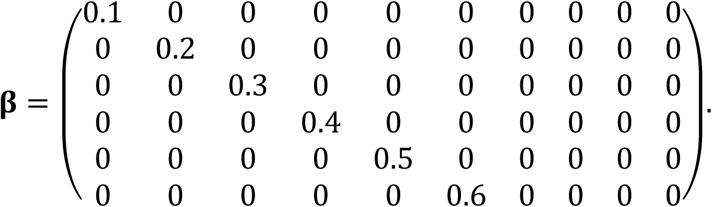

These effect sizes were chosen so that the perturbations explain about 0.2% to 8% of the total variance of each factor.

We generated 300 random datasets under each scenario and performed GFSA analysis. For each dataset, Gibbs sampling was performed for 3000 iterations, and posterior means of parameters were computed from the last 1000 iterations.

We evaluated the results by whether the factors are recovered, and whether the genes affected by a perturbation are identified. Due to the interchangeability of factors in matrix factorization (Equation (1)), we map each of the true factors to the GSFA inferred factor that it is maximally correlated with using the absolute Pearson correlation. The correlations of the true and inferred factors are then assessed. To evaluate the identification of genes affected by perturbations, we define the ground truth, as the genes with non-zero weights on the factors affected by a perturbation.

### Evaluation of differential gene expression (DGE) methods on simulated data

For comparison, we applied Welch’s t-test^28^ to both the normal data and count-normalized deviance residual data. For count data scenarios, we also applied edgeR-QLF^13^ and MAST^29^. In all these DE tests, cells with each perturbation were compared with all other cells without this perturbation for all genes, FDR was computed following the Benjamini-Hochberg procedure for genes under each perturbation, and significant DEGs were obtained by thresholding FDR < 0.05.

### GSFA analysis of CD8^+^ T cell CROP-seq dataset

Raw `cellranger` outputs of the CD8^+^ T cell CROP-seq study^10^ were downloaded from Gene Expression Omnibus (GEO: GSE119450). We merged resting and stimulated T cells from two donors using the R package `Seurat_4.0.1`^49^. We first filtered cells that contain fewer than 500 expressed genes or more than 10% of total read counts from mitochondria genes. Next, we transformed the raw counts into deviance residuals for all genes in all cells, kept the top 6000 genes ranked by deviance statistics (Supplementary Note 2.1), then regressed out unique UMI count, library size and percentage of mitochondrial gene expression from the reduced deviance residual matrix. The resulting matrix was then scaled so that each gene has variance 1.

The gRNA perturbation data are binarized, with gRNAs targeting the same gene deemed as the same type of perturbation. The scaled gene expression matrix and the perturbation matrix were used as input for GSFA. To capture potentially different effects of CRISPR perturbation under resting and stimulated conditions, we used the modified GSFA model with two cell groups (Supplementary Note 3.2), stratifying all cells by their stimulation states (unstimulated:0, stimulated:1). We specified 20 factors in the model. Gibbs sampling was performed for 4000 iterations, and posterior means of parameters were computed from the last 1000 iterations.

We assessed the calibration of the GSFA results using permutation. We created 10 permutation sets on the stimulated and the unstimulated cells, separately. In each permutation set, the cell labels were permuted independently of the perturbation conditions, and GSFA was run on each of these datasets. The calibration was assessed in a few ways. We checked the distribution of PIPs of the perturbation effects on factors **(β)**, and the distribution of LSFRs from the inferred perturbation to gene effects. We expect PIPs to be close to 0 and LSFRs close to 1 in the permutation results. We also assessed the empirical p-values of correlations between perturbations and inferred factors. Since we do not expect any correlation between the two under permutation, any deviation of p-values from the null distribution would indicate that GSFA incorrectly borrows information from perturbations to infer factors, a potential problem that would inflate the results. All permutation results are reported in Fig. S5.

### GSFA analysis of LUHMES CROP-seq dataset

Raw `cellranger` outputs of the LUHMES neural progenitor cell CROP-seq study^37^ were downloaded from Gene Expression Omnibus (GEO: GSE142078). We merged all 3 batches of LUHMES CROP-seq raw data together using the R package `Seurat_4.0.1`^49^, and filtered cells with a library size over 20000 or more than 10% of total read counts from mitochondria genes. Similarly, we transformed the raw count matrix into a reduced deviance residual matrix with top 6000 genes ranked by deviance residual (Supplementary Note 2.1). Differences in experimental batch, unique UMI count, library size, and percentage of mitochondrial gene expression were all regressed out. Running of GSFA is similar to before, except that there is only one cell group, and that Gibbs sampling was run for 3000 iterations. We also assessed calibration of GSFA results, in the same way as we did in the T cell analysis. The results are reported in Fig. S7.

### Running alternative DGE methods on CD8^+^ T cell and LUHMES CROP-seq data

For both stimulated T cells and LUHMES CROP-seq data, we performed alternative DGE analyses for comparison. We applied edgeR-QLF^13^, DESeq2^12^, and MAST^29^ directly to the scRNA-seq raw count data, contrasting cells with each perturbation from those without, for all the genes selected for GSFA. For the LUHMES dataset, experimental batch was included as one of the covariates in these 3 tests. We also applied scMAGeCK-LR^48^ to the transformed and corrected CROP-seq data (described above). For all these methods, FDR was computed following the Benjamini-Hochberg procedure for genes under each perturbation, and significant DEGs were obtained under an FDR cutoff of 0.05.

To assess the calibration of the differential expression test p-values from these methods, we carried out permutation tests for each DGE method by randomly shuffling the cell labels independent of the perturbation conditions. For the T cell dataset, shuffling occurred within the stimulated cells. We generated 10 permuted datasets, and performed the DGE methods in the same way as before.

### Gene ontology enrichment analysis

Gene ontology (GO) over-representation analyses were performed using the `WebGestaltR()` function in the R package `WebGestaltR_0.4.4`^50^ with default parameters and the functional category for enrichment analysis set to the GO Slim “Biological Process” category (`geneontology_Biological_Process_noRedundant`). To interpret GSFA inferred factors (gene modules), genes with weight PIP > 0.95 were treated as the foreground, while all genes used in GSFA were treated as the background in the over-representation analysis. To interpret DEGs discovered under each perturbation by GSFA or other DGE methods, genes with LSFR < 0.05 (or FDR < 0.05) were treated as the foreground, while all genes evaluated were treated as the background in the over-representation analysis.

## Availability

The software implementing GSFA and all source code used in our study have been deposited at https://github.com/gradonion/GSFA and https://github.com/gradonion/GSFA_paper.

## Acknowledgments

We would like to thank N. Gonzales and D. Leach for feedback and revision on the manuscript; M. Stephens for helpful discussions; A. Selewa for help and insights on scRNA-seq data analysis; P. Carbonetto and Y. Liu for assistance with the use of alternative tools. Computing resources were provided by the University of Chicago Research Computing Center. The work was supported by NIH grants R01MH110531, R01HG010773, R01MH116281 (to X.H.), R01 GM126553, R01 HG011883 (to M.C.), and additional grants, NSF 2016307 and Sloan Research Fellowship (to M.C.).

## Author contributions

X.H. conceived the idea. X.H. and M.C. supervised the project. Y.Z. developed the method, implemented the software, and performed the analyses. K.L. tested the software and verified the reported results. Y.Z., X.H. and M.C. wrote the manuscript.

## Supplementary materials

Supplementary figures S1 to S7;

Supplementary tables 1 and 2;

Supplementary note.

