## Supplementary Materials for "A novel Bayesian factor analysis method improves detection of genes and biological processes affected by perturbations in single-cell CRISPR screening"

### Supplementary Figures

#### Figure S1

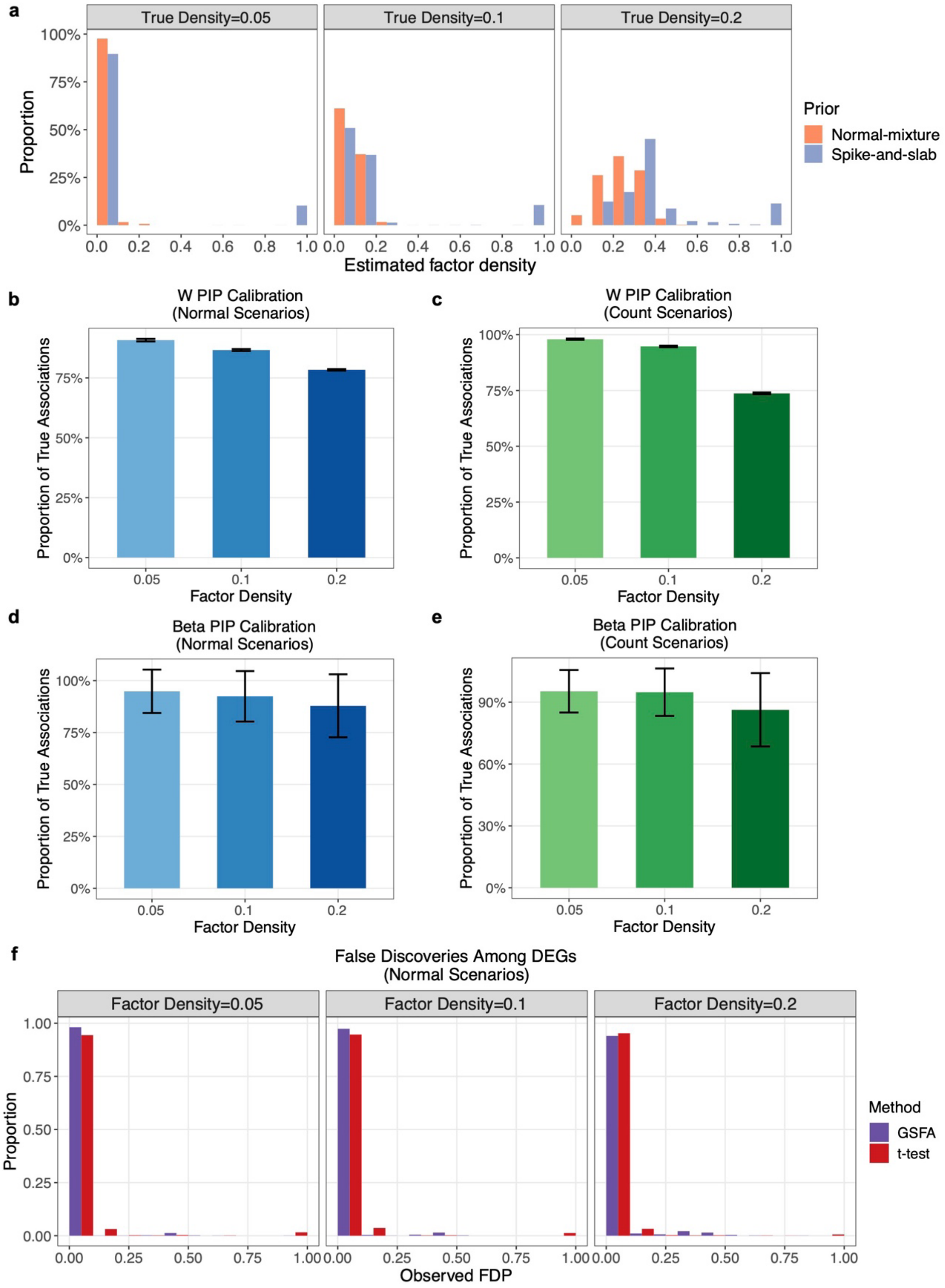

**Figure S1:** Additional GSFA results on simulated data. **a)** Comparison of estimated factor densities using two priors under the count-based setting. **b)** The proportion of truly associated factor-gene pairs out of all the pairs that have GSFA estimated gene loading PIP > 0.95 in the corresponding factor, computed for each dataset under three levels of true factor density and the normal setting. The range of each error bar is one standard deviation ± mean, computed over the proportion values from 300 datasets. **c)** Similar to b) but under the count-based setting. **d)** The proportion of truly associated perturbation-factor pairs out of all the pairs that have GSFA estimated association PIP > 0.95, computed for each dataset under three levels of true factor density and the normal setting. The range of each error bar is one standard deviation ± mean, computed over the proportion values from 300 datasets. **e)** Similar to d) but under the count-based setting. **f)** Observed proportion of false discoveries among significant DEGs detected by GSFA (LFSR < 0.05) or Welch's t-test (FDR < 0.05), computed for each dataset under three levels of true factor density and the normal setting.

#### Figure S2

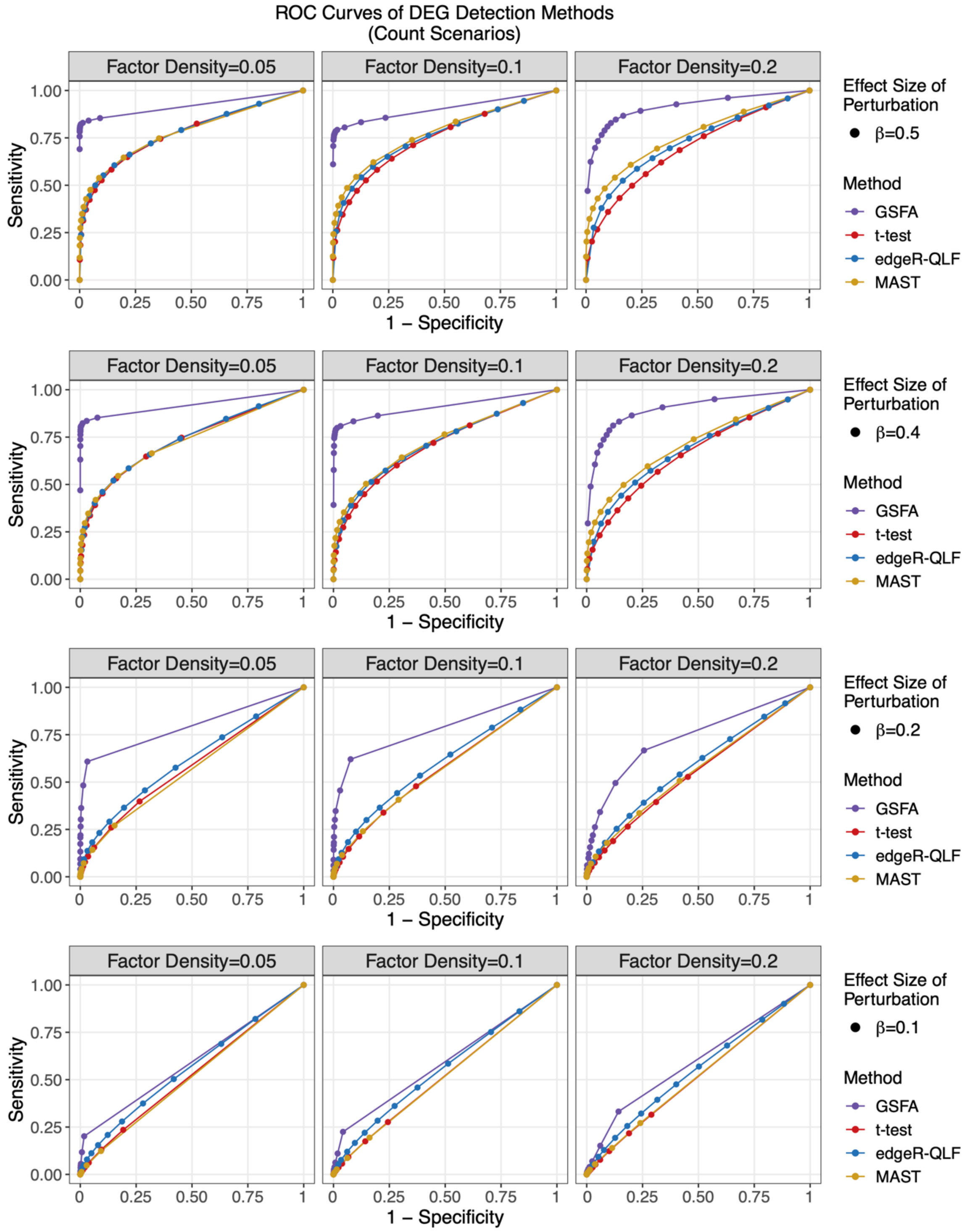

**Figure S2:** ROC curves of DEG discovery across methods on count-based simulated data. Results are across 3 different levels of true factor density, and 4 different values of true perturbation effects; 4 colors correspond to 4 DEG detection methods.

**Figure S3**

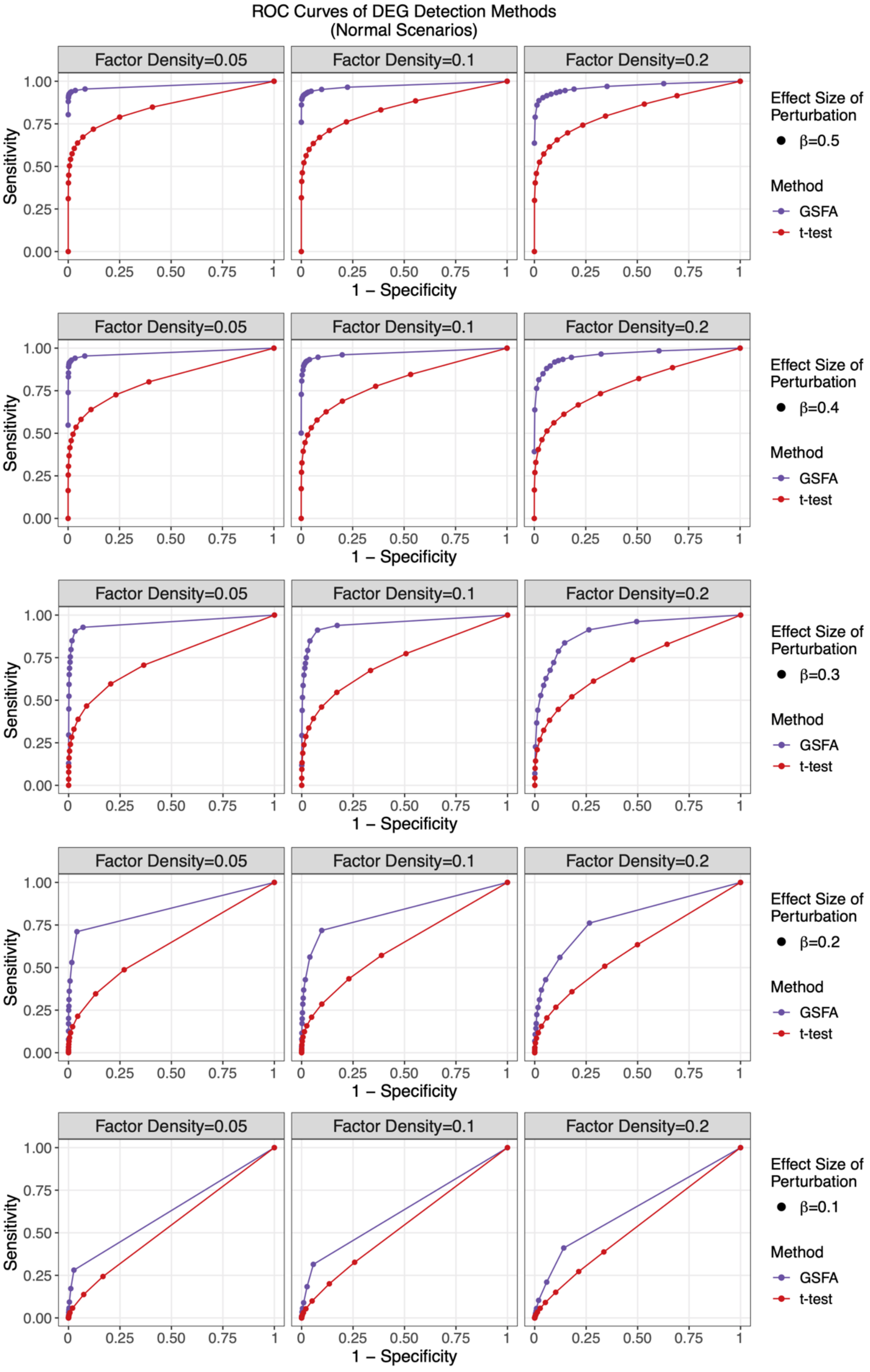
ROC Curves of DEG Detection Methods
(Normal Scenarios)
Factor Density=0.05
Factor Density=0.1
Factor Density=0.2
Sensitivity
1 – Specificity
Effect Size of Perturbation
β=0.5
β=0.4
β=0.3
β=0.2
β=0.1
Method
GSFA
t-test

**Figure S3:** ROC curves of DEG discovery across methods on normal scenario simulated data. Results are across 3 different levels of true factor density, and 5 different values of true perturbation effects; 2 colors correspond to 2 DEG detection methods.

**Figure S4**

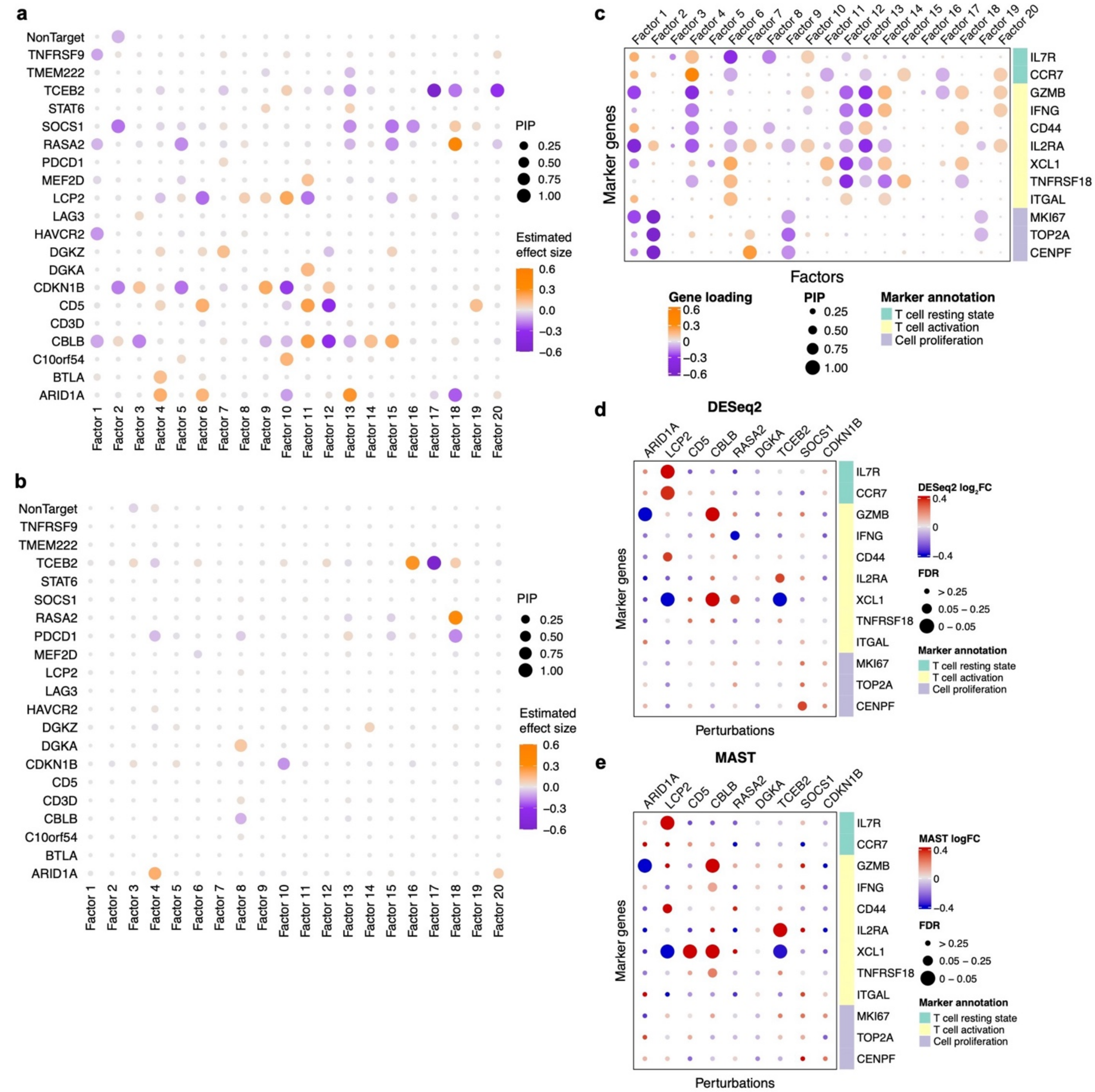

**Figure S4:** Additional GSFA results on CD8$^+$ T cell CROP-seq dataset. **a)** Estimated effects of gene perturbations on all factors inferred by GSFA within stimulated T cells. The size of a dot represents the PIP of association; the color represents the effect size. **b)** Similar to a) but estimated within unstimulated T cells. **c)** Loading of selected marker genes on all factors. The size of a dot represents the gene PIP in a factor and the color represents the gene weight (magnitude of contribution) in a factor. **d)**, **e)** Estimated effects of perturbations on marker genes in stimulated T cells with DESeq2 (d) and MAST (e). Sizes of the dots represent FDR bins; colors of the dots represent the DESeq2 $\log_2$ fold change estimates and the MAST log fold change estimates, respectively.

#### Figure S5

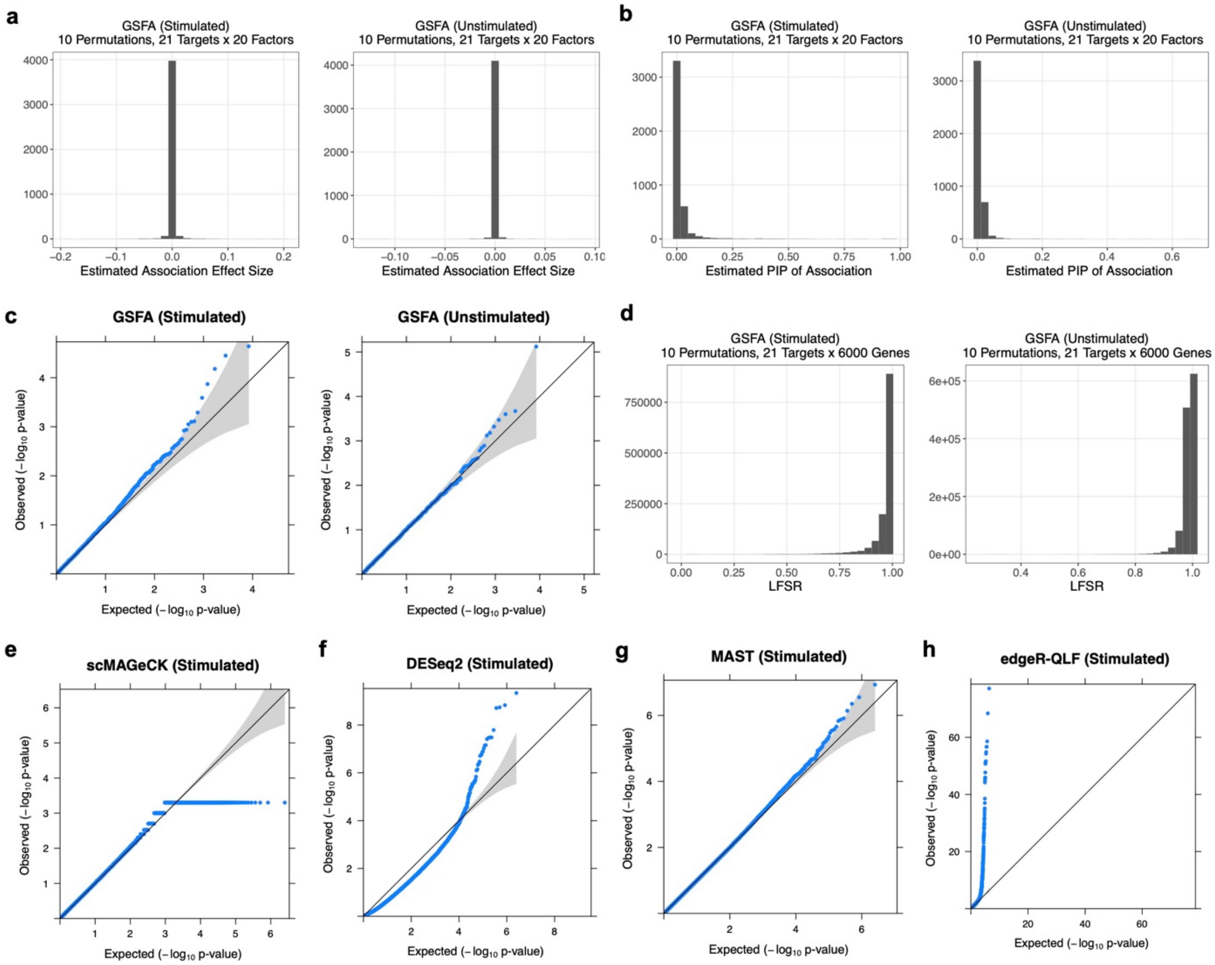

**Figure S5:** Permutation results of DEG detection methods on CD8$^+$ T cell CROP-seq data. Results from 10 randomly permuted datasets are presented together. **a)** GSFA effect sizes of perturbations on factors, estimated within stimulated cells and unstimulated cells, respectively. **b)** GSFA PIPs of associations between factors and perturbations, estimated within stimulated cells and unstimulated cells, respectively. **c)** Q-Q plot of p-values obtained from linear regression between GSFA estimated factors and perturbations within stimulated cells and unstimulated cells, respectively. **d)** GSFA LFSRs of genes under all perturbations, estimated within stimulated cells and unstimulated cells. **e)** Empirical p-values of differential expression estimated by scMAGeCK-LR within stimulated cells; exact zeros were replaced with 5e-4 for visualization in the Q-Q plot. **f)**, **g)**, **h)** Differential expression p-values estimated within stimulated cells by DESeq2 (f), MAST (g), and edgeR-QLF test (h).

### Figure S6

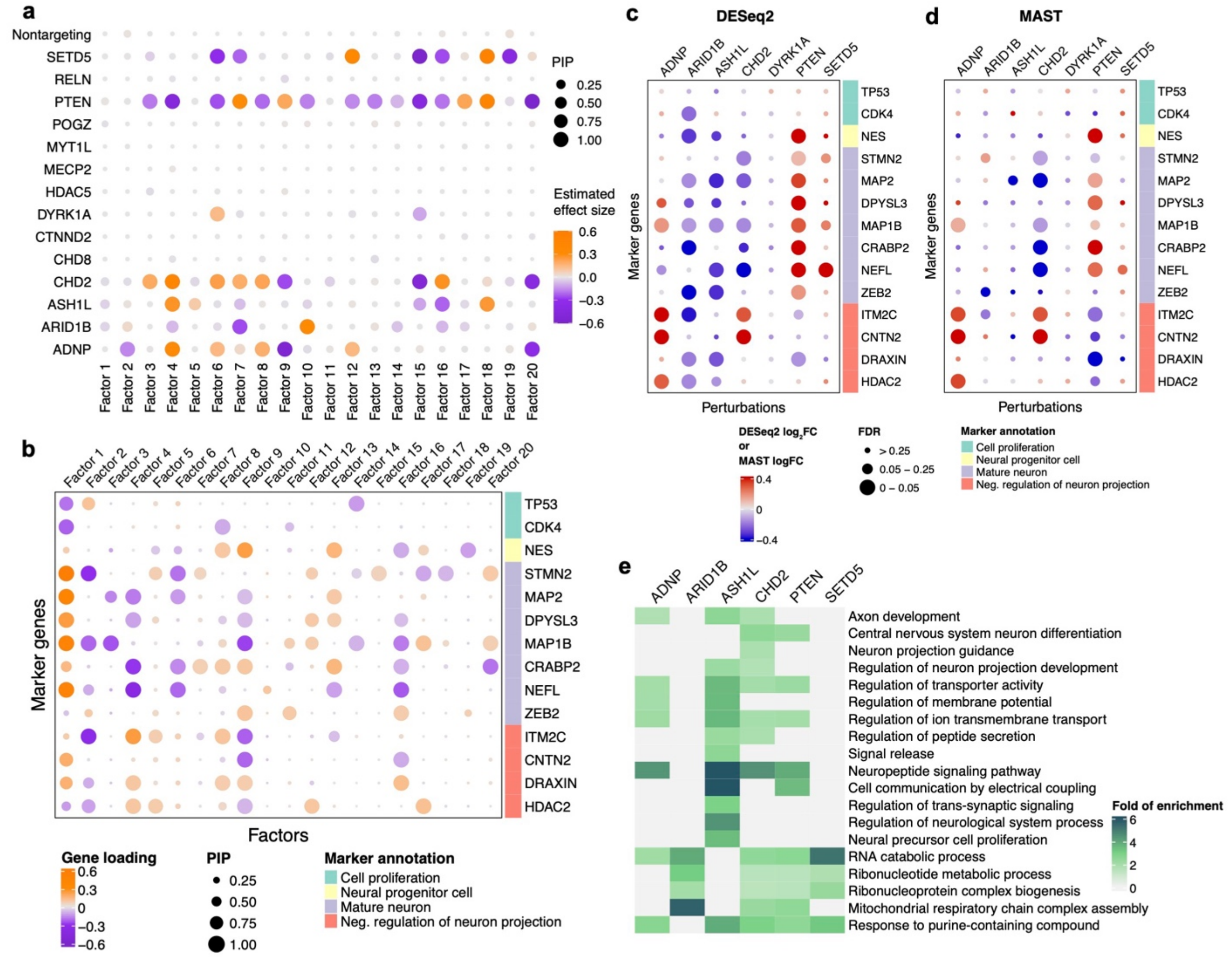

**Figure S6:** Additional GSFA results on LUHMES CROP-seq dataset. **a)** Estimated effects of gene perturbations on all factors inferred by GSFA. The size of a dot represents the PIP of association; the color represents the effect size. **b)** Loading of neuronal marker genes on all factors. The size of a dot represents the gene PIP in a factor and the color represents the gene weight (magnitude of contribution) in a factor. **c)** DESeq2 estimated effects of perturbations on marker genes. Sizes of the dots represent FDR bins; colors of the dots represent the $\log_2$ fold change estimates. **d)** MAST estimated effects of perturbations on marker genes. Sizes of the dots represent FDR bins; colors of the dots represent the log fold change estimates. **e)** Heatmap of selected GO "biological process" terms and their folds of enrichment in DEGs detected by GSFA (LFSR < 0.05).

**Figure S7**

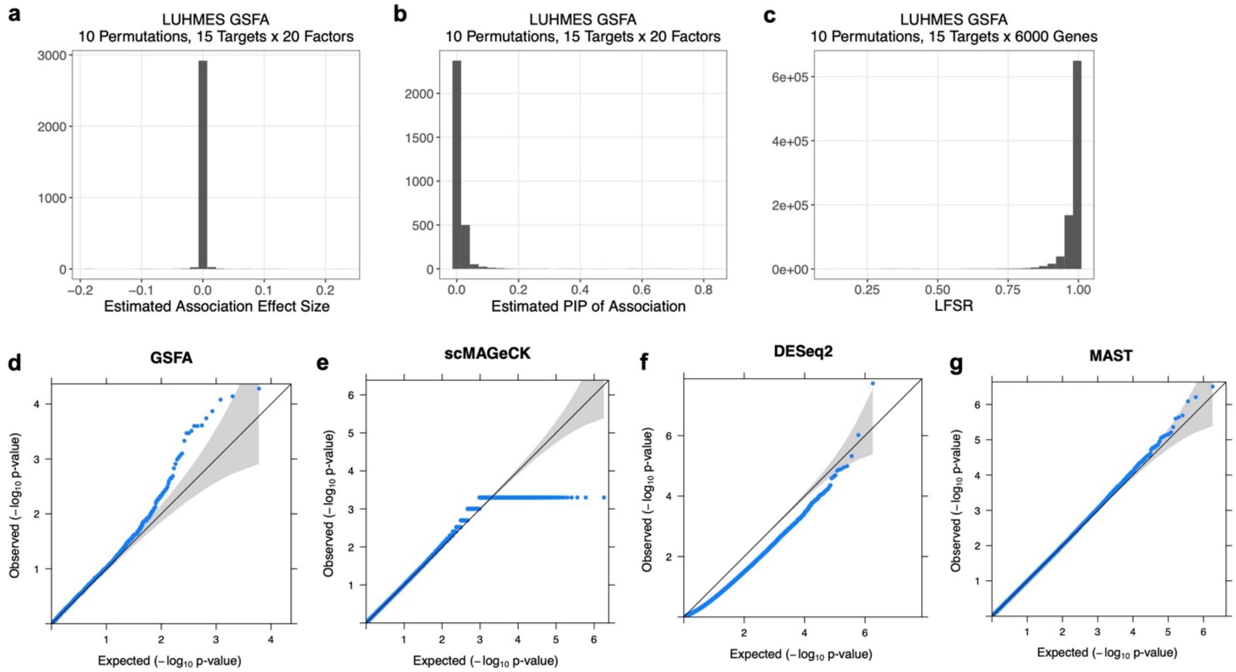

**Figure S7:** Permutation results of DEG detection methods on LUHMES CROP-seq dataset. Results from 10 randomly permuted datasets are presented together. **a)** GSFA effect sizes of perturbations on factors. **b)** GSFA estimated PIPs of associations between factors and perturbations. **c)** GSFA estimated LFSRs of genes under all perturbations. **d)** Q-Q plot of p-values obtained from linear regression between GSFA estimated factors and perturbations. **e)** Empirical p-values of differential expression estimated by scMAGeCK-LR; exact zeros were replaced with 5e-4 for visualization in the Q-Q plot. **f)**, **g)** Differential expression p-values estimated by DESeq2 (f) and MAST (g).

#### Supplementary Tables

##### Table S1. T cell marker genes

| Gene | Protein (Aliases) | Annotation | References |
|---|---|---|---|
| IL7R | Interleukin-7 receptor (CD127) | T cell resting state | PMID: 15308108 |
| CCR7 | CC chemokine receptor 7 | T cell resting state | PMID: 11145663 |
| GZMB | Granzyme B | T cell activation | PMID: 12360212, PMID: 22084442 |
| IFNG | Interferon gamma | T cell activation | PMID: 11145690 |
| CD44 | | T cell activation | PMID: 12526810 |
| IL2RA | Interleukin-2 receptor | T cell activation | PMID: 18417224 |
| XCL1 | X-C motif chemokine ligand 1 | T cell activation | PMID: 19913446 |
| TNFRSF18 | Glucocorticoid-induced TNFR-related protein (GITR) | T cell activation | PMID: 21076066 |
| ITGAL | Integrin subunit alpha L (LFA-1) | T cell activation | PMID: 29774029 |
| MKI67 | Marker of proliferation Ki-67 | Cell proliferation | PMID: 29322240 |
| TOP2A | DNA topoisomerase II alpha | Cell proliferation | PMID: 15980158 |
| CENPF | Centromere protein F | Cell proliferation | PMID: 16565862 |

**Table S2. Neuronal marker genes**

| Gene | Protein (Aliases) | Annotation | References |
|---|---|---|---|
| TP53 | Tumor protein p53 | Cell proliferation | PMID: 18948956 |
| CDK4 | Cyclin dependent kinase 4 | Cell proliferation | PMID: 19733543 |
| NES | Nestin | Neural progenitor cell | PMID: 29541793 |
| STMN2 | Stathmin-2 | Mature neuron | PMID: 14598370 |
| MAP2 | Microtubule associated protein 2 | Mature neuron | PMID: 10704996 |
| DPYSL3 | Dihydropyrimidinase like 3 | Mature neuron | GO:0010976 |
| MAP1B | Microtubule associated protein 1B | Mature neuron | GO:0010976 |
| CRABP2 | Cellular retinoic acid binding protein 2 | Mature neuron | GO:0010976 |
| NEFL | Neurofilament Light Chain | Mature neuron | GO:0010976 |
| ZEB2 | Zinc finger E-box binding homeobox 2 | Mature neuron | GO:0010976 |
| ITM2C | Integral membrane protein 2C | Negative regulation of neuron projection | GO:0010977 |
| CNTN2 | Contactin-2 | Negative regulation of neuron projection | GO:0010975 |
| DRAXIN | Dorsal inhibitory axon guidance protein | Negative regulation of neuron projection | GO:0010977, PMID: 24832731 |
| HDAC2 | Histone deacetylase 2 | Negative regulation of neuron projection | GO:0010977 |
| GO:0010975 regulation of neuron projection development | | | |
| GO:0010976 positive regulation of neuron projection development | | | |
| GO:0010977 negative regulation of neuron projection development | | | |

### SN1 Model specification and inference

#### 1.1 Additional prior specification in GSFA

In line with the Bayesian framework, we specify the following conjugate prior distributions for parameters $\boldsymbol{\psi}$, $\boldsymbol{\pi}$, $\boldsymbol{\sigma}^2$, $\boldsymbol{c}^2$, $\boldsymbol{p}$, and $\boldsymbol{d}^2$ in the GSFA model:

$$\psi_j^{-1} \sim \text{Gamma}(g_0, h_0) \tag{1}$$
$$\pi_k \sim \text{Beta}(s_w r_w, s_w(1 - r_w)) \tag{2}$$
$$\sigma_k^{-2} \sim \text{Gamma}(g_w, h_w) \tag{3}$$
$$c_k^{-2} \sim \text{Gamma}(g_c, h_c) \tag{4}$$
$$p_m \sim \text{Beta}(s_b r_b, s_b(1 - r_b)) \tag{5}$$
$$d_m^{-2} \sim \text{Gamma}(g_b, h_b) \tag{6}$$

In practice, these hyperparameters are set to the following values: $g_0 = 1, h_0 = 1, s_w = 50, r_w = 0.2, g_w = 1, h_w = 1, g_c = 3, h_c = 0.5, r_b = 0.2, g_b = 1, h_b = 1$. In the simulation study, $s_b = 5$; in real data applications, $s_b = 20$.

By choosing these hyperparameters, we set the mean values of the prior distributions of our parameters $\bar{\pi}_k = 0.2$, $\bar{p}_m = 0.2$, $\bar{\psi}_j = \bar{\sigma}_k^2 = \bar{d}_m^2 = 1$, and $\bar{c}_k^2 = 1/6$.

#### 1.2 Gibbs sampling steps in GSFA

Here we describe the details of the Gibbs sampling steps in GSFA.

To obtain posterior samples for $\beta_{mk}$ and $\gamma_{mk}$, we first sample $\gamma_{mk}$ based on the product of two ratios: the ratio of two marginal likelihoods, and the prior ratio.

$$\begin{aligned}\frac{P(\gamma_{mk} = 1|\cdot)}{P(\gamma_{mk} = 0|\cdot)} &= \frac{P(\mathbf{Z}|\gamma_{mk} = 1, \mathbf{G}, \boldsymbol{\beta}_{-mk}, \boldsymbol{\gamma}_{-mk}, \boldsymbol{d}^2)}{P(\mathbf{Z}|\gamma_{mk} = 0, \mathbf{G}, \boldsymbol{\beta}_{-mk}, \boldsymbol{\gamma}_{-mk}, \boldsymbol{d}^2)} \cdot \frac{P(\gamma_{mk} = 1|p_m)}{P(\gamma_{mk} = 0|p_m)} \\ &= \sqrt{\frac{L_{mk}}{d_m^2}} \exp\Big(\frac{\mu_{mk}^2}{2L_{mk}}\Big) \cdot \frac{p_m}{1 - p_m},\end{aligned} \tag{7}$$

where $\mu_{mk} = L_{mk} \sum_{i=1}^N G_{im}(Z_{ik} - \sum_{l:l \neq m} G_{il}\beta_{lk})$ and $L_{mk} = (\sum_{i=1}^N G_{im}^2 + \frac{1}{d_m^2})^{-1}$.

With $\gamma_{mk}$ sampled, we can obtain posterior samples of $\beta_{mk}$ with

$$\beta_{mk}|\gamma_{mk} = 1 \sim N(\mu_{mk}, L_{mk}), \tag{8}$$
$$\beta_{mk}|\gamma_{mk} = 0 \sim \delta_0. \tag{9}$$

For the remaining parameters, we can obtain their posterior samples as follows:

$$W_{j\cdot}|\cdot \sim N((\mathbf{Z}^T\mathbf{Z} + \mathbf{D}_j)^{-1}\mathbf{Z}^T Y_{\cdot j}, \psi_j(\mathbf{Z}^T\mathbf{Z} + \mathbf{D}_j)^{-1}), \tag{10}$$
$$\text{where } \mathbf{D}_j = \text{diag}\Big(\frac{\psi_j}{\sigma_1^2[F_{j1} + (1 - F_{j1})c_1^2]}, ..., \frac{\psi_j}{\sigma_K^2[F_{jK} + (1 - F_{jK})c_K^2]}\Big),$$

$$F_{jk}|\cdot \sim \text{Bern}\Big(\frac{r_{jk}}{r_{jk}+1}\Big), \text{ where } r_{jk} = \frac{\pi_k}{1-\pi_k} c_k \exp\Big[\frac{W_{jk}^2}{2\sigma_k^2}(\frac{1}{c_k^2}-1)\Big], \tag{11}$$

$$Z_{i\cdot}|\cdot \sim N(\mu_i, \Sigma), \tag{12}$$

$$\text{where } \mu_i = \Sigma \cdot (\mathbf{W}^T \mathbf{\Psi}^{-1} Y_{i\cdot} + \boldsymbol{\beta} G_i), \text{ and } \Sigma = (\mathbf{W}^T \mathbf{\Psi}^{-1}\mathbf{W} + \mathbf{I}_K)^{-1},$$

$$\psi_j|\cdot \sim \text{InverseGamma}\Big(g_0 + \frac{N}{2}, h_0 + \frac{1}{2}\sum_{i=1}^{N}(Y_{ij} - \sum_{k=1}^{K} Z_{ik}W_{jk})^2\Big) \tag{13}$$

$$\pi_k|\cdot \sim \text{Beta}(s_w r_w + \sum_{j=1}^{P} F_{jk}, s_w(1-r_w) + P - \sum_{j=1}^{P} F_{jk}) \tag{14}$$

$$\sigma_k^2|\cdot \sim \text{InverseGamma}\Big(g_w + \frac{P}{2}, h_w + \frac{1}{2}\sum_{j=1}^{P} \frac{W_{jk}^2}{F_{jk} + (1-F_{jk})c_k^2}\Big) \tag{15}$$

$$c_k^2|\cdot \sim \text{InverseGamma}\Big(g_c + \frac{1}{2}\sum_{j=1}^{P}(1-F_{jk}), h_c + \frac{1}{2}\sum_{j:F_{jk}=0} \frac{W_{jk}^2}{\sigma_k^2}\Big) \tag{16}$$

$$p_m|\cdot \sim \text{Beta}(s_b r_b + \sum_{k=1}^{K} \gamma_{mk}, s_b(1-r_b) + K - \sum_{k=1}^{K} \gamma_{mk}) \tag{17}$$

$$d_m^2|\cdot \sim \text{InverseGamma}\Big(g_b + \frac{1}{2}\sum_{k=1}^{K}\gamma_{mk}, h_b + \frac{1}{2}\sum_{1=k}^{K} \beta_{mk}^2\Big) \tag{18}$$

In practice, $\mathbf{Z}$ and $\mathbf{W}$ are initialized from a truncated singular value decomposition (SVD) of the normalized gene expression matrix $\mathbf{Y}$, with the number of left (and right) singular vectors being $K$, the number of factors specified in the model. The last 20% quantile of elements (in terms of absolute value) in $\mathbf{W}$ are set to 0, and $\mathbf{F}$ is initialized as the binarized version of $\mathbf{W}$. $\beta_m$ is initialized as the coefficients of linear regression $\mathbf{Z} \sim G_m$ $(1 \leq m \leq M)$. The last 50% quantile of elements (in terms of absolute value) in $\boldsymbol{\beta}$ are set to 0, and $\boldsymbol{\gamma}$ is initialized as the binarized version of $\boldsymbol{\beta}$.

The initialization of additional parameters is as follows: $\psi_j^{(0)} = 1$, $\pi_k^{(0)} = 0.2$, $\sigma_k^{2(0)} = 1$, $c_k^{2(0)} = 0.25$, $p_m^{(0)} = 0.2$, $d_m^{2(0)} = 1$ $(1 \leq j \leq P$, $1 \leq k \leq K$, $1 \leq m \leq M)$.

### SN2 Input pre-processing

#### 2.1 Deviance residual transformation and feature selection for count data

To accommodate the application of GSFA on count data, we follows the transformation proposed in [1], where the count data are transformed into continuous quantities in the form of deviance residuals. In a standard data normalization pipeline, raw counts are normalized by sample-specific size factors, and then log-transformed. However, due to the large number of zeros in scRNA-seq UMI counts, normalization schemes commonly used for bulk RNA-seq data may result in unstable normalization [2], and the arbitrary pseudocount added during the log transformation of exact zeros may introduce systematic errors and cause spurious differences in expression [3]. The deviance residual transformation circumvents these difficulties by directly modeling the raw count data under a multinomial null model of constant gene expression across all cells, and quantifying the fit of data in the form of deviance residuals, a quantity analogous to z-scores and approximately follow a normal distribution. Specifically, the deviance residual for gene $j$ in cell $i$ is

$$r_{ij} = \text{sign}(c_{ij} - \hat{\mu}_{ij})\sqrt{2c_{ij}\log\frac{c_{ij}}{\hat{\mu}_{ij}} + 2(n_i - c_{ij})\log\frac{n_i - c_{ij}}{n_i - \hat{\mu}_{ij}}}. \tag{19}$$

Here $c_{ij}$ is the raw gene count, $n_i$ is the library size of cell $i$, and $\hat{\mu}_{ij} = n_i \frac{\sum_i c_{ij}}{\sum_i n_i}$ is the expression of gene $j$ under the null model of constant expression.
Following [1], we use an approximate multinomial deviance statistic to evaluate the deviance of a gene from the null model:

$$D_j = \sum_{i=1}^{N} r_{ij}^2. \tag{20}$$

Genes with constant expression across cells are not informative and will have a deviance of 0, while genes that vary across cells in expression will have a larger deviance. Therefore, one can pick the genes with high deviance during feature selection as an alternative to selecting highly variable genes, with the advantage that the selection is not sensitive to normalization.

### SN3 Alternative models

#### 3.1 GSFA model with alternative prior on gene weights

GSFA also allows one to use the standard spike-and-slab prior for the gene weights, although we find that it does not work as well as the default mixture-of-normal prior (Equation (4) in Methods). The alternative spike-and-slab prior is given by:

$$W_{jk} \sim \pi_k N(0, \sigma_k^2) + (1 - \pi_k)\delta_0. \tag{21}$$

Similarly, we introduced a latent binary matrix $\mathbf{F}_{P\times K}$ to indicate whether $W_{jk}$'s are nonzero. We can obtain the posterior samples of these parameters as follows:

$$\frac{P(F_{jk} = 1|\cdot)}{P(F_{jk} = 0|\cdot)} = \sqrt{\frac{\lambda_{jk}}{\sigma_k^2}}\,\exp\Big(\frac{\nu_{jk}^2}{2\lambda_{jk}}\Big)\cdot\frac{\pi_k}{1 - \pi_k}, \tag{22}$$

where $\nu_{jk} = \lambda_{jk}\sum_{i=1}^{N} Z_{ik}(Y_{ij} - \sum_{h:h\neq k} Z_{ih}W_{jh})/\psi_j$ and $\lambda_{jk} = (\sum_{i=1}^{N} Z_{ik}^2/\psi_j + 1/\sigma_k^2)^{-1}$.

With $F_{jk}$ sampled, we can obtain posterior samples of $W_{jk}$ with

$$W_{jk}|F_{jk} = 1 \sim N(\nu_{jk}, \lambda_{jk}), \tag{23}$$
$$W_{jk}|F_{jk} = 0 \sim \delta_0. \tag{24}$$

#### 3.2 GSFA model with multiple cell groups

In the cases when one is interested in learning about the effects of perturbations under different cell types or experimental conditions, we extend the current model (Equations (1) - (4) in Methods) so that the factors are inferred using all cells but the associations between factors and perturbations are estimated separately for each cell group. For example, assuming 2 groups of cells, group 0 and group 1, we have the conditional probability of factor matrix $\mathbf{Z}$

$$P(\mathbf{Z}|\mathbf{G}, \boldsymbol{\beta_0}, \boldsymbol{\beta_1}) = \prod_{i0 \in \text{group } 0} N(Z_{i0\cdot}; \boldsymbol{\beta_0} G_{i0\cdot}, \mathbf{I}_K) \prod_{i1 \in \text{group } 1} N(Z_{i1\cdot}; \boldsymbol{\beta_1} G_{i1\cdot}, \mathbf{I}_K), \tag{25}$$

where $\boldsymbol{\beta_0}$ and $\boldsymbol{\beta_1}$ are $M \times K$ matrices holding the effect sizes of perturbations on factors within group 0 cells and group 1 cells, respectively.
Each effect size matrix is still subjected to the same "spike-and-slab" prior:

$$\beta_{0mk} \sim p_{0m} N(0, d_{0m}^2) + (1 - p_{0m})\delta_0, \tag{26}$$

$$\beta_{1mk} \sim p_{1m} N(0, d_{1m}^2) + (1 - p_{1m})\delta_0. \tag{27}$$

The distributions of other model parameters remain the same.

Once we have the posterior samples of parameters and latent variables, we can similarly obtain the posterior samples of the total effects of perturbations on individual genes, $\theta_{mj}$'s, within each cell group using Equation (5) in Methods and the corresponding $\beta_{mk}$ for that cell group.
